# Motif-driven microRNA regulation of B cell tolerance uncovers ERα as a female-specific checkpoint

**DOI:** 10.1101/2025.01.22.634242

**Authors:** Jaime L Díaz-Varela, Beatriz Herrero-Fernández, María del Pilar González-Molina, Laura Gámez-Reche, Ana María Prieto-Muñoz, Javier Sanz, Marina Mendieta-Homs, MariPaz López-Molina, Tania Gonzalo, Alba Moreno, Laura Mañas, David Nemazee, Changchun Xiao, Laura Otero-Ortega, Alicia González-Martín

## Abstract

Breakdown of B cell tolerance is a central feature of autoimmune diseases, yet the molecular mechanisms underlying the female predominance of these diseases remain unclear. Here, we identified miR-130b as an essential component of a miRNA network, defined by the GUGCA seed motif, that regulates central B cell tolerance. Elevated miR-130b levels impaired tolerance by promoting the survival of immature B cells upon B cell antigen receptor engagement. This network converged on the estrogen receptor pathway through downregulation of *Esr1* and *Pten*. Genetic ablation of *Esr1* in immature B cells was sufficient to compromise central B cell tolerance in females, but not in males, revealing an unrecognized role for ERα in this process. In males, *Esr1* deficiency reduced bone marrow B cell numbers, whereas in females, B cell numbers were preserved, with a greater proportion of cells displaying lower surface CD19 levels. Transcriptomic analysis revealed sex-specific genetic programs characterized by defective B cell development and skewed immunoglobulin gene usage in males, and increased PI3K-AKT signaling in females. In females, this molecular rewiring compensated for developmental defects, but also attenuated central tolerance mechanisms, allowing the escape of autoreactive B cells to the periphery. In patients with multiple sclerosis, elevated levels of miR-130b in circulating vesicles correlated with more severe disease, including increased formation of new demyelinating lesions, cognitive decline, and neurodegeneration. Together, our findings identify a sex-specific ERα-dependent checkpoint in B cell tolerance controlled by a seed-driven miRNA network, providing a mechanistic framework for female predisposition to autoimmunity.

## Main

Autoimmune diseases comprise more than 80 chronic, and often disabling, pathologies caused by dysregulated immune responses against self-tissues. Together, they affect 8-10% of the population (*1, 2*) and, for reasons that are poorly understood, their incidence is rising (*2, 3*). Most autoimmune diseases remain incurable, and current therapies are frequently associated with significant adverse effects and high socioeconomic costs. Notably, these pathologies are markedly more prevalent in females than males (*4*). However, the cellular and molecular mechanisms underlying this female predominance are incompletely understood.

Autoimmune diseases are characterized by a breakdown of immune tolerance, the series of mechanisms that prevent lymphocytes from attacking self-tissues while generating a large receptor repertoire capable of eliminating a large variety of invading pathogens and arising tumors. Autoreactive B cells play critical roles in autoimmunity through the generation of autoantibodies, presentation of autoantigens to T cells and secretion of proinflammatory cytokines (*5*). Previous studies showed that B cell tolerance is defective in patients with autoimmune diseases including multiple sclerosis, lupus erythematosus and rheumatoid arthritis (*6–8*), but the molecular mechanisms that controls it remain understudied.

B cell tolerance is initiated in the bone marrow. During early stages of B cell development, genetic rearrangements, known as V(D)J recombination, occur to form the immunoglobulin (Ig) heavy and light chains, respectively, resulting in an IgM molecule with a given specificity. Upon expression of IgM on the cell surface, the B cell enters the immature stage. The stochastic nature of V(D)J recombination promotes diversification of the B cell receptor repertoire, enabling recognition of a broad range of foreign antigens, but also generates autoreactive specificities. To prevent the exit of potentially harmful autoreactive B cells to the periphery, immature B cells are subjected to the central tolerance checkpoint. During this process, non-autoreactive cells exit the bone marrow to mature in the periphery, while autoreactive cells undergo Ig light chain rearrangement to acquire a new antigen specificity through receptor editing. If this process eliminates autoreactivity, the cell proceeds to peripheral maturation and, otherwise, it can undergo additional rounds of receptor editing. If this mechanism fails to eliminate self-reactivity, the cells are removed from the repertoire by clonal deletion (apoptosis) (*9*). Additional tolerance checkpoints exist in the periphery, referred to as peripheral B cell tolerance, to regulate autoreactive B cells that have escaped central tolerance, including anergy, a state of non-responsiveness (*10*).

The IgM^b^-macroself mouse model provides a robust *in vivo* platform for identifying novel regulators of B cell tolerance (*11*). These mice ubiquitously express an engineered superantigen that recognizes the constant region of the IgM heavy chain. In this model, early B cell development occurs normally and, upon IgM expression by immature B cells, binding to the superantigen occurs, mimicking self-recognition. This results in the elimination of all immature B cells through clonal deletion and, consequently, the absence of mature B cells in peripheral sites, namely the spleen and lymph nodes. Using this model, we previously established microRNAs (miRNAs) as key regulators of B cell tolerance, specifically miR-148a and miR-19b (*12, 13*). However, the identified mechanisms did not explain the increased susceptibility of women to autoimmune diseases compared to men. In this study, through the identification of a motif-defined miRNA network, we uncovered Estrogen receptor 1 (*Esr1*) as essential for maintaining adequate control of central B cell tolerance in females only, revealing a previously unknown function for ERα that may contribute to the higher prevalence of autoimmune diseases in women.

### Identification of a minimal sequence expressed in miRNAs that regulate B cell tolerance

MiRNAs are ∼22 nucleotide non-coding RNAs that regulate gene expression by binding the 3′ untranslated regions (UTRs) of target mRNAs, leading to translational repression or mRNA degradation (*14*). Target recognition relies mainly on nucleotides 2-8, known as the seed sequence, and miRNAs sharing the same seed are grouped into families. Previous studies showed that miRNAs regulate many processes of immune cell development and function (*15*), and that the expression of many miRNAs is dysregulated in patients with autoimmune diseases (*16*).

To uncover functionally relevant miRNA networks in B cells, we performed *in silico* analysis on the seed sequence regions (positions 1–8) of 101 lymphocyte-expressed miRNAs (Extended Data Table 1), reasoning that sequence patterns that occur in a group of related sequences often correlate with a biological function (Fig. 1a). This analysis identified multiple 4-6 nucleotide consensus motifs, one of which, GUGC, was highly conserved across multiple miRNA families, suggesting functional relevance. Eight miRNA families, miR-455, miR-33, miR-210, miR-19, miR-17, miR-148, miR-147 and miR-130, contained this motif (Fig. 1b). We then examined the expression levels of these families in B cells during development and upon activation. Most of them were expressed at moderate to high levels and, within families, the member with the highest expression levels was chosen for further analysis (Extended Data Fig.1a). This resulted in six individual miRNAs: miR-33a-5p, miR-210-3p, miR-19b-3p, miR-17-5p, miR-148a-3p and miR-130b-3p (referred to as miR-33a, miR-210, miR-19b, miR-17, miR-148a and miR-130b hereafter) (Fig. 1c). MiR-147 and miR-455 were undetectable or expressed marginally (if at all), respectively, in immature B cells and were consequently excluded from the study (Extended Data Fig.1b).

**Fig. 1.**
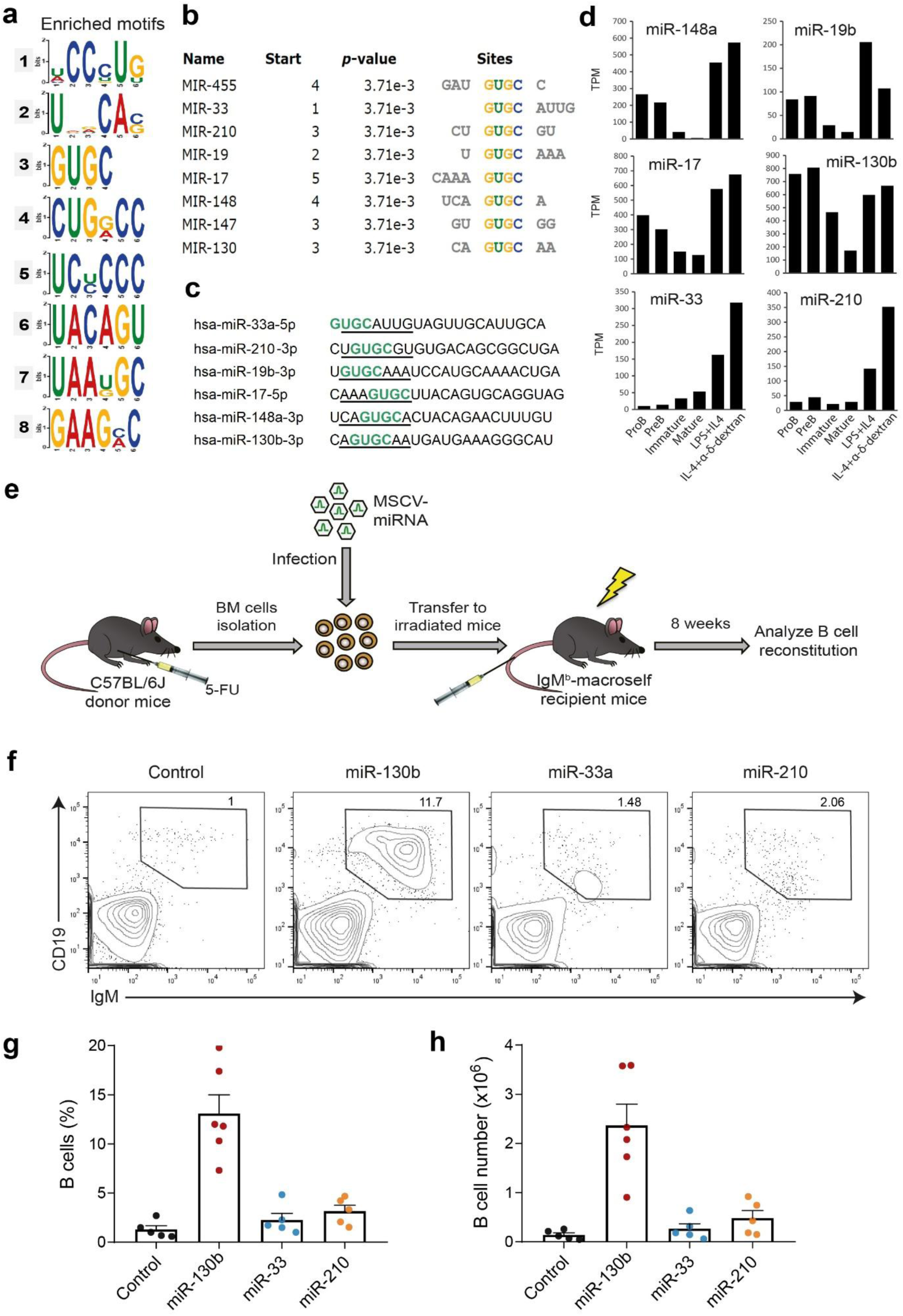
MiR-130b promotes escape of B cells from central B cell tolerance. **a,** Hit motifs from Multiple Em for Motif Elicitation (MEME) analysis of 101 lymphocyte-expressed miRNA families. Nucleotides 1-8 of each miRNA, that contain their seed sequence, were used as input for the analysis. **b,** MiRNA families containing the conserved GUGC motif, ranked third by MEME. **c,** Representative miRNA members within each family with the highest expression levels in B lymphocytes. The seed regions are underlined, and the GUGC motif highlighted in green. **d,** Expression levels of miRNAs in **c** throughout murine B cell development and activation as indicated. The graphs in **d** were generated based on previously published data (*33*). **e,** Schematic representation of bone marrow transduction and reconstitution experiments in the murine B cell tolerance model IgM^b^-macroself. **f,** Representative flow cytometry contour plots showing splenic B cells of IgM^b^-macroself mice reconstituted with bone marrow transduced with miR-130b, miR-33, miR-210 or control vector, as indicated. Numbers above the outlined area indicate percent of CD19^+^-IgM^+^ cells (splenic B cells) among total splenocytes. **g,h,** Bar graph showing the percentages (**g**) and total numbers (**h**) of splenic B cells of all mice analyzed. Graphs represent mean + s.e.m. Data are pooled from five independent experiments in **g**,**h**. n=5-6 mice/group.

This candidate list included miR-148a and miR-19b, the only miRNAs previously shown to regulate B cell tolerance (*12, 13*), suggesting that additional miRNAs within this group might control this process. In addition, the expression levels of miR-17 and miR-130b throughout B cell development and activation displayed a similar pattern to that of miR-148a and miR-19b. Specifically, these miRNAs were expressed in proB and preB cells, downregulated in immature B cells, further downregulated in mature B cells in the periphery, and induced upon activation (Fig. 1d). In contrast, miR-33 and miR-210 followed different expression patterns. Of note, the potential role of miR-17, a family member of the miR-17-92 cluster, in B cell tolerance was already analyzed in our previous work, which excluded a major role for this miRNA in B cell tolerance (*13*).

### miR-130b regulates B cell tolerance

To validate the potential role of the identified miRNAs in the regulation of B cell tolerance, a hematopoietic stem and precursor cell (HSPC) transduction and reconstitution approach was performed in the mouse model of clonal deletion IgM^b^-macroself, as previously described (*12*). Briefly, bone marrow (BM) cells from donor C57BL/6J mice inoculated with 5-fluorouracil 4 days before were isolated, enriched for HSPCs, transduced with MSCV-based retroviral particles encoding miR-130b, miR-33, miR-210 or an empty construct (control), and used to reconstitute lethally irradiated IgM^b^-macroself mice for 8 weeks (Fig. 1e). This system provides a clean readout, since the presence of B cells in the spleen of these mice after reconstitution indicates a breakdown of B cell tolerance mediated by the transduced miRNA.

Analysis of the spleens of IgM^b^-macroself mice after reconstitution revealed that increased levels of miR-130b promote the escape of a significant percentage of B cells from clonal deletion and their exit from the bone marrow to the periphery, with approximately 12% of splenic B cells among total splenocytes (Fig. 1f-h). Conversely, miR-33a or miR-210 did not mediate any substantial effect in this process, as the percentages of splenic B cells were close to those detected in the spleens of IgM^b^-macroself mice reconstituted with control-transduced HSPCs, which had approximately 1% of B cells among total splenocytes (*12, 13*). These results demonstrate a function for miR-130b in the regulation of B cell tolerance. In line with this, elevated serum levels of miR-130b have been associated with renal damage in patients with lupus nephritis (*17*).

### Enrichment of predicted target genes in Estrogen receptor signaling

To identify novel regulators of B cell tolerance we built upon the observation that all lymphocyte-expressed miRNAs containing the minimal sequence GUGCA within their seed, miR-130b, miR-148a and miR-19b, control this process. We hypothesized that they regulate B cell tolerance through key common signaling pathways. The predicted targets of these miRNAs were retrieved from miRDB and, as shown in the Venn diagram, 110 targets were common to the three miRNAs (Fig. 2a and Extended Data Table 2). These targets were then subjected to Ingenuity Pathway Analysis (IPA) to identify signaling pathways highly enriched in proteins that may play a regulatory role in central B cell tolerance. Estrogen Receptor Signaling was retrieved as the top-ranked canonical pathway (Fig. 2b,c). Eight predicted targets containing binding sites for these miRNAs formed part of this pathway: *Cfl2*, *Esr1*, *Igf1*, *Med12L*, *Prkaa1*, *Pten*, *Runx2* and *Sos2*.

**Fig. 2.**
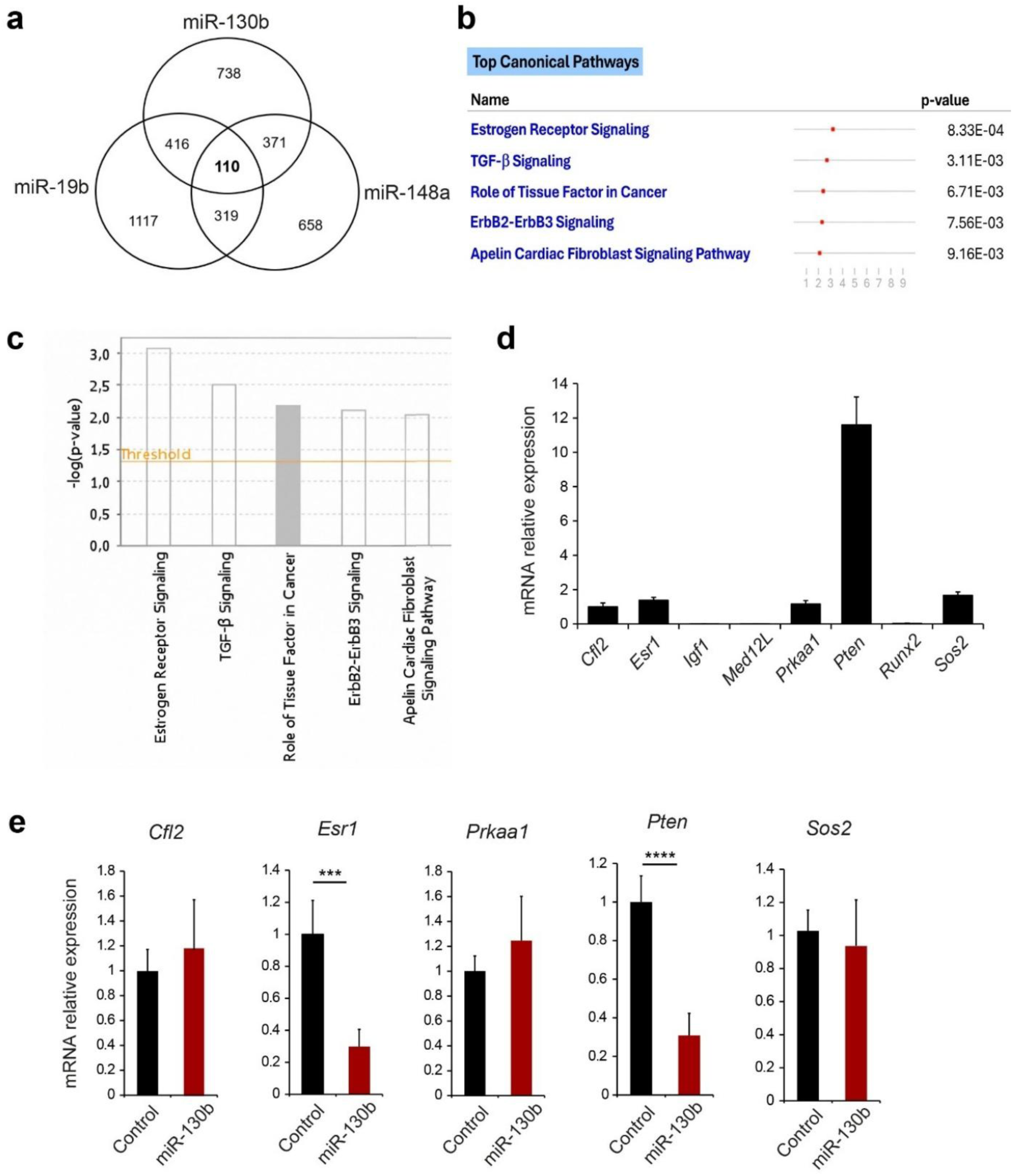
The Estrogen receptor signaling pathway is enriched in targets regulated by miRNAs that control B cell tolerance. **a,** Venn diagram of the predicted target genes (miRDB) of miR-130b, miR-19b and miR-148a, the miRNAs identified as regulators of B cell tolerance to date. **b,c,** Top canonical pathways identified through Ingenuity Pathway Analysis (IPA) of the 110 common target genes of these miRNAs. The orange line in **c** represents the threshold for a positive z-score. **d,** Bar graph showing the relative expression of all the target genes included in the estrogen receptor signaling pathway, as indicated, in primary murine immature B cells sorted from C57BL/6J mice and analyzed by qRT-PCR. Graphs show mean + s.d. in immature B cells sorted from bone marrow of 19 mice. **e,** Bar graphs showing relative expression of *Cfl2*, *Esr1*, *Prkaa1*, *Pten* and *Sos2* in WEHI-control and WEHI-miR-130b cells stimulated for 14 hours. Data were pooled from two independent experiments in **e**. Statistical significance was determined with an unpaired two-tailed Student’s t test. *** P ≤ 0.001 and **** P ≤ 0.0001.

Since miRNA regulation is cell-type dependent, we next determined whether these proteins are expressed in immature B cells. Immature B cells from bone marrow of C57BL/6J mice were stained with antibodies against cell markers CD19, CD93 and IgM coupled with fluorophores and purified through fluorescence-activated cell sorting (FACS). This was followed by extraction of total RNA from these samples and analysis of the expression levels of these targets using quantitative real-time polymerase chain reaction (qRT-PCR). *Cfl2*, *Esr1*, *Prkaa1*, *Pten* and *Sos2* exhibited moderate to high expression levels in immature B lymphocytes (Fig. 2d). In contrast, *Igf1*, *Med12L* and *Runx2* displayed marginal or negligible expression and were therefore excluded from further analyses.

As the targetomes of miRNAs are also cell context-dependent (*15*), we assessed whether these targets are regulated by miR-130b in the immature B cell line WEHI-231. This cell line recapitulates the clonal deletion process when its IgM molecule is bound with an anti-IgM antibody, mimicking self-recognition and resulting in a fraction of cells undergoing BCR-engagement-induced apoptosis (clonal deletion) as it occurs during central tolerance. Stable WEHI-231 cells expressing or not increased levels of this miRNA were generated through transduction with control- or miR-130b-expressing retroviruses that also encode GFP, followed by enrichment by FACS. WEHI-miR-130b cells showed more robust miR-130b expression compared with WEHI-control cells, as validated through TaqMan assays (Extended Data Fig. 2a).

We next stimulated both stable WEHI-231 cell lines with anti-IgM for 14 hours, a time in which cells are signaling for apoptosis (clonal deletion) but still have not initiated the death process. Cells were then harvested, and total RNA was purified to measure the expression levels of these target genes by qRT-PCR. Among the selected targets, *Esr1* (which encodes for Estrogen receptor alpha; ERα) and *Pten* (which encodes the tumor suppressor PTEN) were downregulated by 70% in cells with increased levels of miR-130b compared to control cells (Fig. 2e). The levels of the other genes were not downregulated by miR-130b.

### ERα functions as a sex-specific regulator of central B cell tolerance

To validate whether these targets regulate B cell tolerance, we obtained mice deficient in *Esr1* and *Pten* and used their bone marrow cells to reconstitute lethally irradiated IgM^b^-macroself mice (Fig.3a). After 8 weeks, the spleens of these mice, as well as those of a group reconstituted with cells from control mice, were analyzed for the escape of B cells to the periphery.

The role of PTEN in B cell tolerance had been previously established by us and others (*12, 13, 18*). When bone marrow cells from donor mice with B cell-specific deletion of *Pten* (CD19Cre;*Pten*^fl/fl^) were used to reconstitute irradiated IgM^b^-macroself mice, a drastic percentage of B cells was observed in the spleens of those mice 8 weeks later compared with the control group reconstituted with bone marrow cells from C57BL/6J wild type mice, which had almost no B cells in their spleens (Fig.3b-d).

The potential role of *Esr1* in central B cell tolerance had not been previously analyzed. Since ERα is expressed and functions differentially in women and men, we specifically addressed the role of *Esr1* in females by reconstituting irradiated IgM^b^-macroself mice with bone marrow cells from *Esr1*-deficient female mice (*Esr1* KO). As expected, mice in the control group had a nearly complete absence of B cells in their spleens. In contrast, deficiency of *Esr1* was sufficient to enable escape from B cell tolerance, as indicated by the presence of approximately 14% of B cells in the spleens of IgM^b^-macroself mice reconstituted with bone marrow cells from *Esr1^-/-^* mice (Fig. 3e-g). These results demonstrate that ERα is required for the adequate establishment of central B cell tolerance, a previously unknown function for this receptor. Next, we asked whether this effect was sex-specific. To address this, the same experimental approach was performed using both female and male mice. Interestingly, the requirement for ERα for B cell tolerance was observed in female, but not male, mice (Fig. 3h-j), uncovering a novel sex-dependent mechanism promoting autoimmunity.

**Fig. 3.**
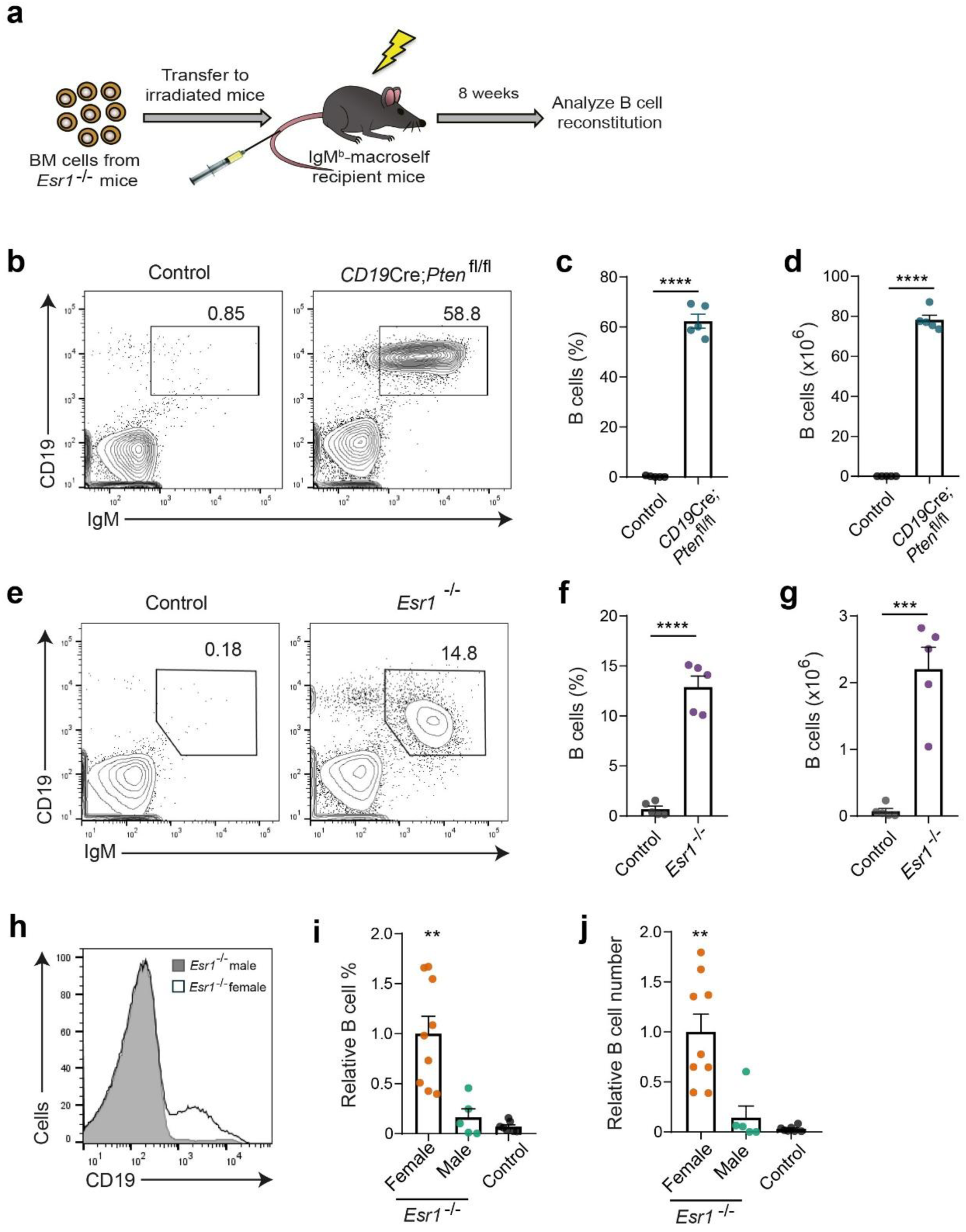
ERα deficiency impairs B cell tolerance in a sex-dependent manner. **a,** Outline showing bone marrow reconstitution experiments of IgM^b^-macroself mice. **b,e,** Representative contour plots showing splenic CD19^+^-IgM^+^ B cells in recipient IgM^b^-macroself mice reconstituted with bone marrow (BM) cells from donor *Esr1^-/-^, CD19*Cre*;Pten*^fl/fl^ or control mice, as indicated, analyzed by flow cytometry 8 weeks after reconstitution. Mice reconstituted with *Esr1*^-/-^ BM cells were all female. **c,d,f,g,** Bar graphs showing the percentages (**c,f**) and total numbers (**d,g**) of splenic B cells of all mice analyzed in **b** and **e**. **h,** Representative overlay histogram showing percentages of CD19^+^ B cells in female and male IgM^b^-macroself mice reconstituted with BM cells from *Esr1^-/-^* KO mice. **i,j,** Bar graphs showing relative CD19^+^-IgM^+^ B cells percentages (**i**) and numbers (**j**) in all mice analyzed in **h**, and pooled male and female IgM^b^-macroself mice reconstituted with BM cells from control mice. Female values were arbitrarily set to 1 and all other conditions are shown relative to this group. Graphs represent mean + s.e.m. Data are representative of two independent experiments. n=7 mice (control group), n=5 (*CD19*Cre;*Pten*^fl/fl^ and male *Esr1*^-/-^ groups) and n=9 mice (female *Esr1*^-/-^ group). Statistical significance was determined with an unpaired two-tailed Student’s t test. ** P ≤ 0.01, *** P ≤ 0.001 and **** P ≤ 0.0001.

### *Esr1* deficiency impairs B cell development in male but not female mice

To gain further insight into this sex-specific function of ERα, we examined the B cell compartment in the bone marrow of female and male control and *Esr1^-/-^* mice. Male and female control mice displayed similar B cell percentages and numbers in the bone marrow. *Esr1* deficiency resulted in a significant reduction of B cells in male mice. This decrease, however, was not recapitulated in *Esr1^-/-^* female mice, which displayed similar B cell percentages and numbers as the control groups (Fig. 4a-d and Extended Data Fig. 3a-c). A greater proportion of B cells in these mice displayed reduced surface levels of CD19, a coreceptor that lowers the signaling threshold required for BCR-induced clonal deletion (Fig. 4a,e) (*19*). The proportions of precursor, immature and mature B cells within bone marrow cells remained constant across all male and female control or *Esr1^-/-^*groups of mice, as determined by staining with the markers CD93, IgM and CD19 (Extended Data Fig. 4a-b).

**Fig. 4.**
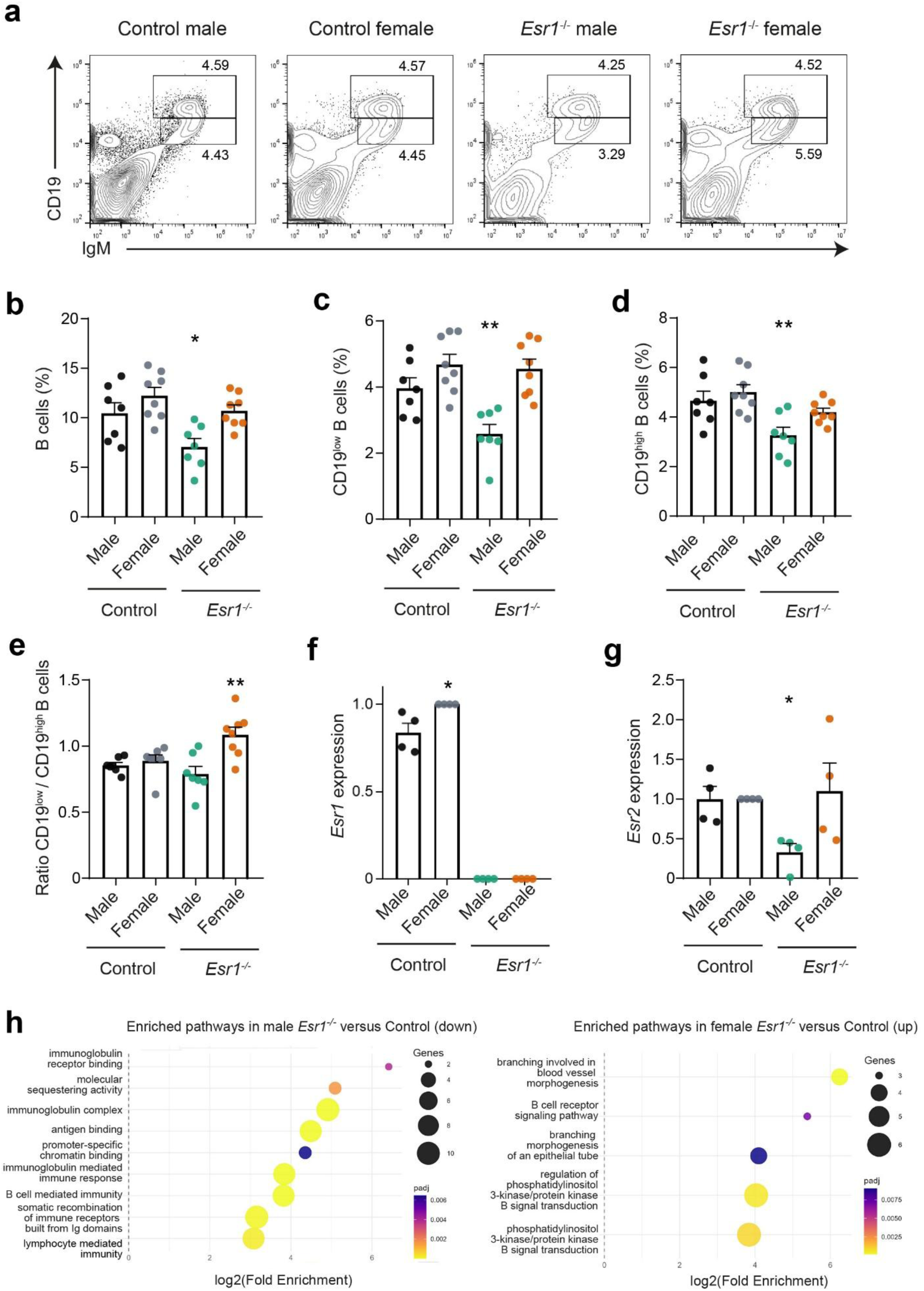
***Esr1*-deficient cells reduce their surface CD19 expression levels in female, but not male, mice. a,** Representative contour plots showing percentages of CD19^low^ and CD19^high^ B cells (CD19^+^-IgM^+^) in the bone marrow of control and *Esr*1^-/-^ male and female mice, as indicated. **b-d.** Bar graphs showing percentages of CD19^+^-IgM^+^ (**b**), CD19^low^-IgM^+^ (**c**) and CD19^high^-IgM^+^ (**d**) cells in the bone marrow of all mice analyzed in **a**. **e,** Graph showing the ratio of CD19^low^ versus CD19^high^ B cells of the mice in **c-d**. **f,g.** Relative mRNA levels of *Esr1* (**f**) and *Esr2* (**g**) in bone marrow B cells from male and female control and *Esr1^-/-^* mice determined by qRT-PCR. **h.** GO enrichment analysis showing pathways enriched in genes downregulated in developing B cells in the bone marrow of male *Esr1^-/-^* mice compared with their control counterparts, or upregulated in female *Esr1^-/-^* cells compared with the control group. Graphs represent mean + s.e.m. Data are pooled from three independent experiments in **a-d** and four in **f-g**. n=7 mice (control and Esr1^-/-^ female groups), and n=8 mice (control and Esr1^-/-^ male groups). Statistical significance was determined with unpaired two-tailed Student’s t tests. * P ≤ 0.05 and ** P ≤ 0.01.

We next measured the expression levels of *Esr1* and *Esr2*, which exhibit partially redundant roles, in developing B cells from the bone marrow of these groups of mice. Purified B cells from female control mice expressed higher levels of *Esr1* than those of males and, as expected, its expression was absent in both *Esr1^-/-^* groups (Fig. 4f). *Esr2* levels are lower than those of *Esr1* in these cells (Extended Data Fig. 3d-e), and their levels were drastically reduced in male *Esr1*-deficient mice compared with control and *Esr1^-/-^*female mice (Fig. 4g).

Our data revealed a male-specific B cell developmental defect in *Esr1*-deficient mice that is rescued in female mice, most likely through an extensive rewiring of signaling networks in precursor and immature B cells. To examine this, we performed transcriptomic analysis of developing B cells from male and female control and *Esr1^-/-^* mice. Indeed, bone marrow B cells from male and female mice showed drastically different transcriptomes in the absence of *Esr1* (Extended Data Fig. 4a,b). The *Esr1^-/-^* male B cell transcriptome was enriched in altered usage of immunoglobulin gene segments and antigen binding pathways, with 12 downregulated (Ighg2c, Igkv1-122, Ighv1-78, Igkv12-41, Ighg3, Ighv1-4, Igkv8-21, Ighv8-12, Ighv1-69, Ighv2-2, Ighv3-1, Ighv3-6) and 5 upregulated (Ighv6-6, Igkv2-112, Ighv5-6, Ighv5-9, and Ighv6-3) Ig fragments (Fig. 4h and Extended Data Fig. 4c,e). These defects are unlikely to be compensated for by *Esr2*, since its levels are drastically reduced in developing B cells from these mice. In contrast, the *Esr1^-/-^* female B cell transcriptome was characterized by increased PI3K-AKT signaling (Cxcl12, Bank1, Igfbp5, Serpine2, Angpt1, and Tek), with CXCL12 as a central node (Fig. 4h and Extended Data Fig. 4c,e). These sex-dependent differences were not present between male and female control mice. Overall, our results indicate that B cell development is compromised in *Esr1^-/-^*males, whereas females compensate this effect through extensive rewiring of their transcriptome to potentiate prosurvival signals through the PI3K-AKT pathway, and increase the threshold for apoptosis through CD19 downregulation. This, in turn, impairs clonal deletion of autoreactive B cells, enabling their escape to the periphery.

### *Esr1* and *Pten* are regulated by miR-130b

To demonstrate whether miR-130b directly regulates ERα and PTEN, we used the immature B cell lines WEHI-231 expressing either increased or basal levels of miR-130b. We confirmed that, while miR-130b has no effect on apoptosis in unstimulated cells, it largely protects these cells from BCR engagement-induced apoptosis upon stimulation with anti-IgM (Fig. 5a,b). No miR-130b-dependent difference in cell proliferation was observed (Fig. 5c). Next, reporter assays were performed to confirm whether miR-130b regulates *Esr1* and *Pten* through direct binding to their cognate binding sites in the 3’UTR regions (Extended Data Table 3). These assays showed that miR-130b reduces the expression levels of the reporter gene fused to the 3’UTRs of both target genes (Fig. 5d). This effect was completely or significantly lost for *Esr1* and *Pten*, respectively, when the binding sites of the miRNA were mutated.

**Fig. 5.**
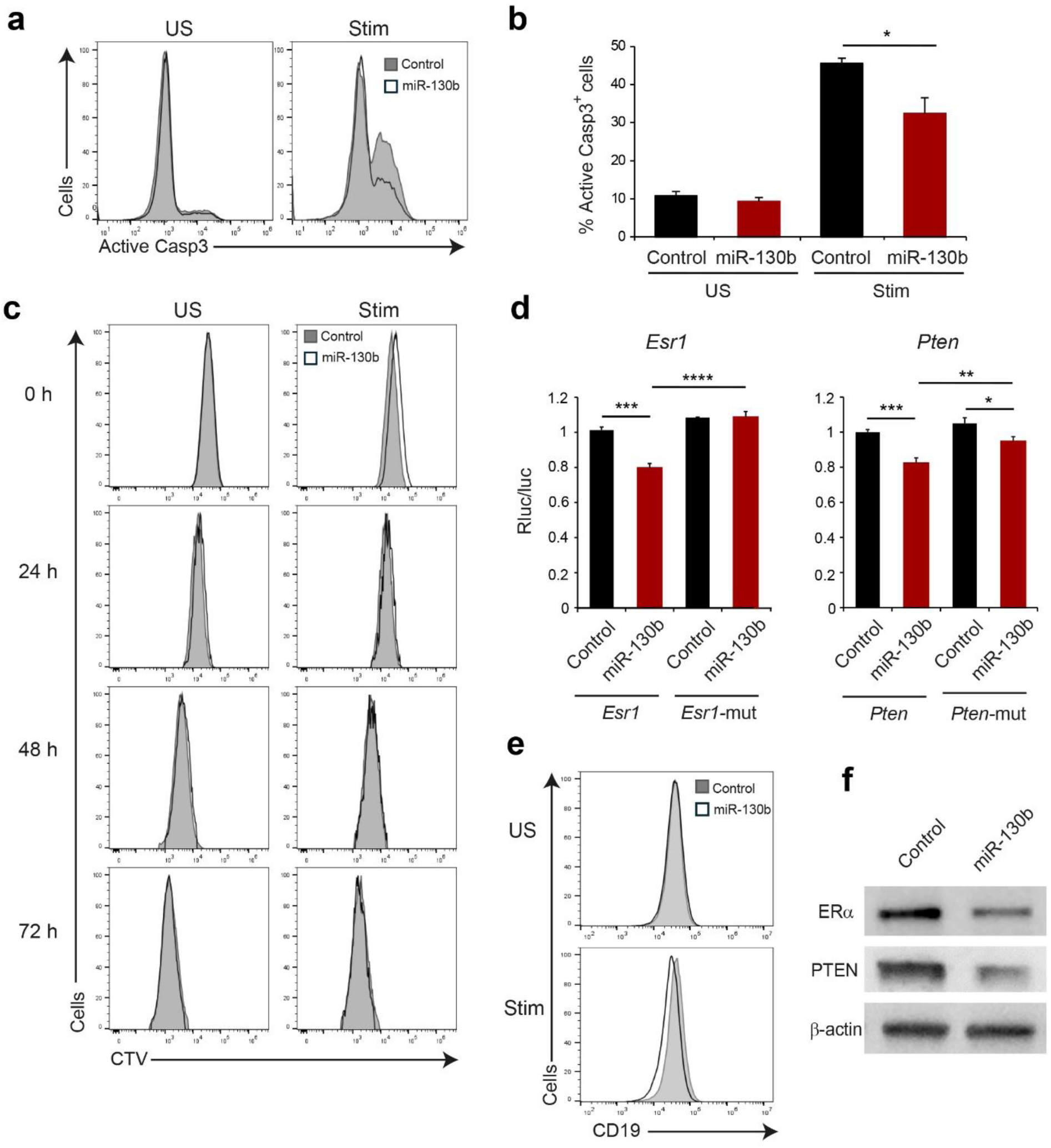
MiR-130b targets *Esr1* and *Pten*. **a,** Representative overlay histograms showing active caspase 3 (Casp3^+^) analysis of WEHI-control and WEHI-miR-130b cells left unstimulated (US) or stimulated (Stim) with anti-IgM (2μg/ml) for 72 hours. **b,** Frequency of active Casp3^+^ cells among cells as in **a**. **c,** Flow cytometry analyzing cells as of **a** labeled with Cell Trace Violet (CTV) at 0, 24, 48 and 72 h. **d,** Bar graphs showing normalized expression of renilla luciferase reporter protein fused to the wild-type 3′ UTR of *Esr1* or *Pten* and 3′ UTR of *Esr1* or *Pten* with mutated binding sites for miR-130b, cotransfected with an expression vector encoding miR-130b or an empty vector (control). The levels of the reporter protein were analyzed 24 hours post-transfection and normalized to that of a constitutively expressed firefly luciferase. **e,** Overlay histograms showing mean fluorescence intensity of CD19 levels in WEHI-control and WEHI-miR-130b cells under unstimulated or stimulated conditions at 48h. **f,** Immunoblot analysis of protein levels of ERα and PTEN, as well as β-actin (loading control), in these cells. Graphs show mean + s.d. Data are representative of two independent experiments in **a**, **c**, **d, e** and **f**, and pooled from two independent experiments in **b**. Statistical significance was determined with an unpaired two-tailed Student’s t test. * P ≤ 0.05, ** P ≤ 0.01, *** P ≤ 0.001 and **** P ≤ 0.0001.

We next measured the levels of membrane-bound CD19 in unstimulated and stimulated conditions in these cells. Consistent with our previous results, surface CD19 levels were downregulated in stimulated conditions in cells with increased miR-130b levels compared with their control counterpart, which aligns with the impaired clonal deletion observed (Fig. 5e). In addition, the protein levels of ERα and PTEN were determined in these cell lines by Western blot. These results confirmed the miR-130b-mediated reduction of ERα and PTEN levels (Fig. 5f). Overall, our data show that both ERα and PTEN expression are reduced by miR-130b through its binding to their cognate binding sites in their 3’UTRs, which increases the BCR signaling threshold required for clonal deletion and, therefore, impairs the elimination of autoreactive B cells.

### Increased serum levels of miR-130b in patients with autoimmunity associate with worse disease activity and severity

Given that B cells are a major source of serum extracellular vesicles (EVs) (*20, 21*), we investigated whether the levels of miR-130b in EVs could serve as a biomarker for patients with autoimmunity. EVs are easier and more cost-effective to isolate from serum than B cells, and they also protect miRNAs from degradation, ensuring more stable and reproducible detection of circulating miR-130b. Neither ERα nor PTEN are present within EVs.

To assess this, serum samples were collected from a cohort of patients with multiple sclerosis (MS; Extended Data Table 4), followed by EV isolation and quantification of miR-130b by NGS. Our analysis showed that elevated levels of miR-130b were associated with more severe disease activity, as indicated by increased number of new lesions in the central nervous system (Fig. 6a). This was accompanied by accelerated cognitive decline and brain atrophy, evidenced by reduced volumes of gray matter, cortical gray matter, total brain, caudate nucleus, and cerebellum (Fig. 6b-g).

**Fig. 6.**
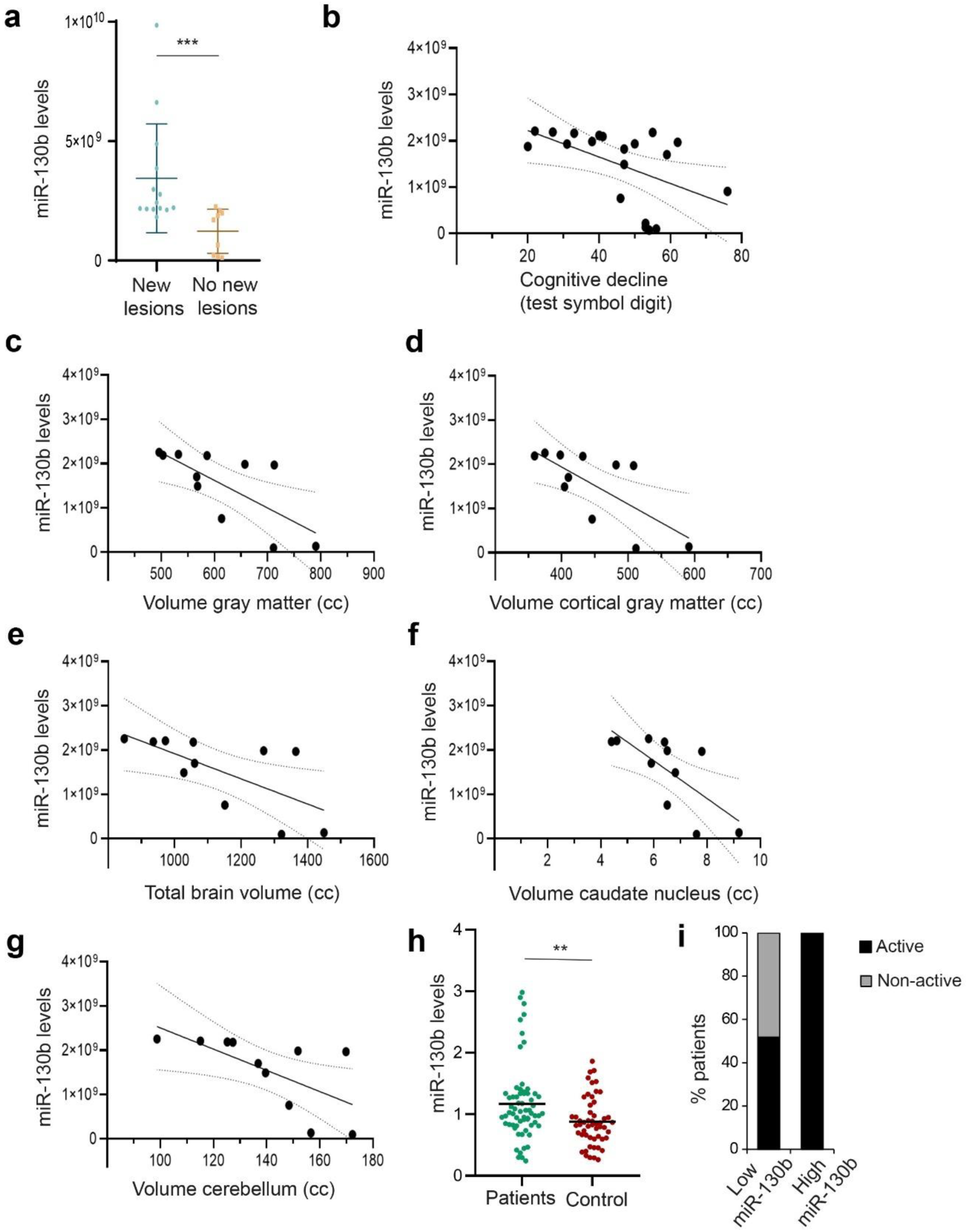
Elevated miR-130b in circulating extracellular vesicles associated with increased disease activity and brain atrophy in patients with multiple sclerosis. **a,** Levels of miR-130b in circulating extracellular vesicles (EVs) of patients with multiple sclerosis developing or not new central nervous system lesions, as determined by magnetic resonance imaging. **b-g,** Cognitive decline (**b**) and volume of gray matter (**c**), cortical gray matter (**d**), total brain (**e**), caudate nucleus (**f**) and total cerebellum (**g**), as a function of miR-130b expression. **h,** Levels of miR-130b in circulating EVs of patients with multiple sclerosis or healthy individuals determined by TaqMan analysis. **i,** Percentages of patients with active disease in the subgroup with higher levels (miR-130b≥2) of miR-130b (12.5%) compared with all other patients (miR-130b<2). Statistical significance was determined with Spearmańs rank test (n= 41 patients in **a**, n=64 and 53 patients and healthy individuals, respectively, in **h, i**). ** P ≤ 0.01, *** P ≤ 0.001.

We next determined whether the levels of miR-130b are higher in all patients with MS compared with healthy individuals. Analysis of their circulating EVs revealed significantly higher relative levels of miR-130b in patients than in healthy controls (Fig. 6h). This difference was partially driven by a subgroup of 8 patients, representing 12.5% of the cohort, who exhibited markedly elevated levels of this miRNA. Analysis of the clinical characteristics of this subgroup showed that all patients were undergoing active disease at the time of sample collection (Fig. 6i), in contrast to the heterogeneous disease states observed in the rest of the cohort, consistent with our previous findings.

In addition, elevated levels of miR-130b in patients correlated with increased percentages of circulating HVEM positive memory B cells, and CD40 positive plasma cells, indicating B cell involvement in disease pathogenesis (Extended Data Fig. 5). Overall, the levels of miR-130b in serum EVs correlated with increased disease activity in patients with multiple sclerosis and, therefore, could serve as a potential biomarker.

## Discussion

Humoral immunity is fundamental not only in adults, but also for fetuses and newborns, who receive antibodies from the mother to protect them from infections during pregnancy and in their first months after birth. Female immunity is known to be generally stronger than that of males. Epidemiological studies have revealed that females are more resistant to infections, showing decreased mortality compared with men and stronger antiviral immune responses (*22*). Females are also more prone to many autoimmune diseases, with a male to female incidence ratio of approximately 1:8 for lupus erythematosus, and 1:3 for rheumatoid arthritis and multiple sclerosis (*4*). Despite extensive efforts, the molecular mechanisms underlying female predisposition to autoimmunity remain incompletely understood.

In this study, we identified estrogen receptor signaling as a key pathway that controls B cell tolerance downstream of miRNAs that regulate this process, defined by a GUGCA motif within their seeds. Within this pathway, ERα, a hormone receptor with markedly different expression levels and functions in females than in males (*23*), and PTEN were validated as central regulators of B cell tolerance. While ERα is a key regulator of reproductive system development and function, including during pregnancy, it also plays important roles in non-reproductive tissues, such as the immune system (*24*). We found that ERα, which is expressed by B cells, is required for adequate B cell development in male mice, since its deficiency resulted in developmental defects and fewer B cells in their bone marrow. This observation is consistent with those from Thurmon et al, 2000, who reported a significant reduction across all bone marrow B lymphocyte subpopulations in male *Esr1* knockout mice, with no significant changes in the hematopoietic progenitor compartment (*25*). However, this study did not examine female mice. Our results uncovered that this developmental defect is compensated in ERα-deficient females by the promotion of prosurvival signals and the elevation of the threshold for BCR engagement-induced apoptosis (clonal deletion) during the immature B cell stage.

From an evolutionary perspective, B cell immunity is particularly important during pregnancy and lactation to provide antibody-mediated protection to fetuses and newborns before they develop their functional immune systems. Indeed, both mice and humans exhibit a shift in the CD4 T cell compartment to a Th2, B cell-supporting subset during pregnancy (*25*). While any given B cell developmental defects in men might be largely compensated by innate and cellular immunity, the same defects in women could be more detrimental because humoral immunity is essential for fetal and neonatal protection through maternal antibody transfer. Our data suggest that B cell development in females is largely rewired at the molecular level to compensate for these defects and, therefore, preserve the generation of the B cell repertoire. However, to achieve this, the threshold for BCR-induced apoptosis increases in immature B cells, which comes at the cost of impaired clonal deletion of autoreactive B cells during central B cell tolerance. These cells escape to the periphery where they could contribute to the onset of autoimmune diseases.

Indeed, ERα has been previously shown to prevent autoimmunity. *Esr1* germline knockout mice developed spontaneous autoimmune glomerulonephritis with formation of splenic germinal centers, proteinuria, infiltration of B cells in the kidneys, damage of tubular cells and serum anti-dsDNA antibodies by one year of age (*26*). Another study partially attributed this autoimmune phenotype to an ERα-mediated control of follicular helper T cell responses (*27*). Here, we provide genetic evidence demonstrating that ERα is required to control autoreactive B cells from exiting the bone marrow to the periphery where they could damage self-tissues in female, but not male, mice. Loss of *Esr1* compromised the central tolerance checkpoint, most likely contributing to the autoimmune disease that developed in *Esr1*^−/−^ mice. In humans, ERα levels were reported to be significantly reduced in freshly isolated T lymphocytes from patients with systemic lupus erythematosus compared to healthy controls (*28*). Concurrently, lower estrogen levels in postmenopausal women have been associated with increased susceptibility to autoimmune diseases (*24*).

Developing B cells from female *Esr1^-/-^* mice showed increased PI3K-AKT signaling compared with those from wild-type females. Downregulation of PTEN, the other identified target of miR-130b within the estrogen receptor pathway, further promotes this prosurvival signal. Indeed, we and others previously showed that PTEN regulates B cell tolerance and autoimmunity (*12, 18, 29*). Studies on the reproductive system showed that the levels of PTEN fluctuate across the menstrual cycle, suggesting its regulation by hormones in the endometrium. 17β-estradiol (E2) stimulates both genomic and nongenomic ERα actions in this system, and the protein levels of PTEN are the highest in uterine epithelial cells during the E2-driven phase of the menstrual cycle (*30*). Another study showed that E2 signaling increases the levels of PTEN and promotes the formation of cytosolic ERα-PTEN complexes in an E2-dependent manner (*31*). This is consistent with functional cooperation between these targets to establish a critical sex-specific checkpoint in central B cell tolerance.

Since miRNAs are frequently contained within circulating extracellular vesicles in humans (*32*), we investigated whether miR-130b in EVs could serve as a biomarker for autoimmune diseases. EVs protect miRNAs from degradation, enabling stable detection of molecular signals in circulation and offering a readily accessible, cost-effective alternative to direct cell-based analyses for clinical applications. Notably, increased miR-130b levels in EVs from patients with multiple sclerosis correlated with more severe disease progression, characterized by the formation of new brain lesions, and accelerated cognitive decline and neurodegeneration. These findings suggest that elevated miR-130b levels in EVs may reflect increased autoreactive activity and potentially serve as a biomarker of disease severity in multiple sclerosis.

Overall, our study identified a sex-dependent ERα mechanism as a critical regulator of B cell tolerance and autoimmunity. These findings advance our current understanding of B cell tolerance and provide new insights into the molecular mechanisms underlying sex differences in autoimmune diseases, with potential diagnostic and therapeutic implications for patients with these diseases.

## Extended data

**Extended Data Fig. 1.**
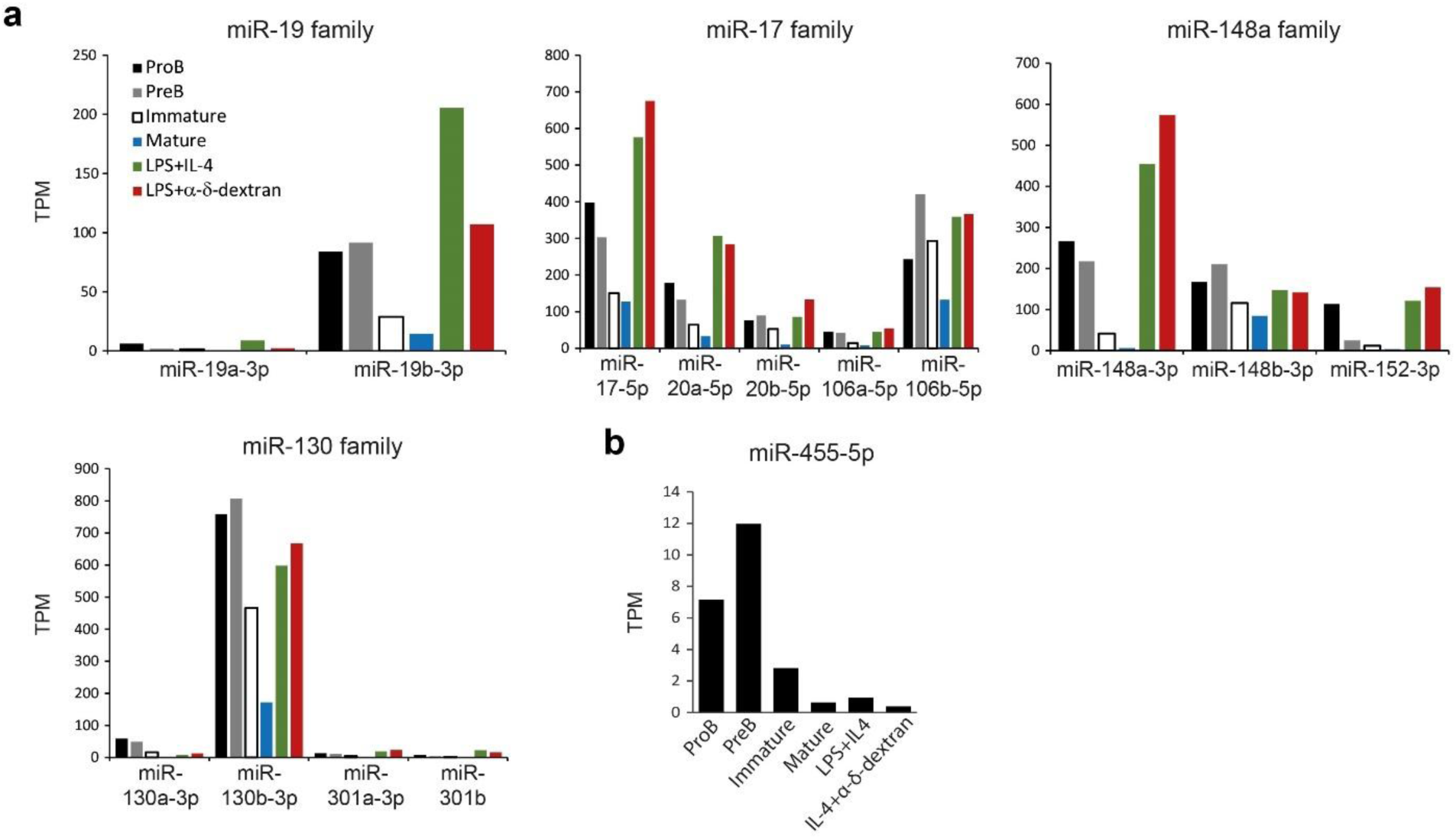
Expression levels of miRNA families identified by MEME analysis and miR-455-5p. **a,b,** Bar graphs showing expression levels of individual miRNAs within the miRNA families identified by MEME analysis (**a**) and miR-455-5p (**b**) at different stages of B cell development and activation, as indicated. The graph was generated based on previously published data (*33*).

**Extended Data Fig. 2.**
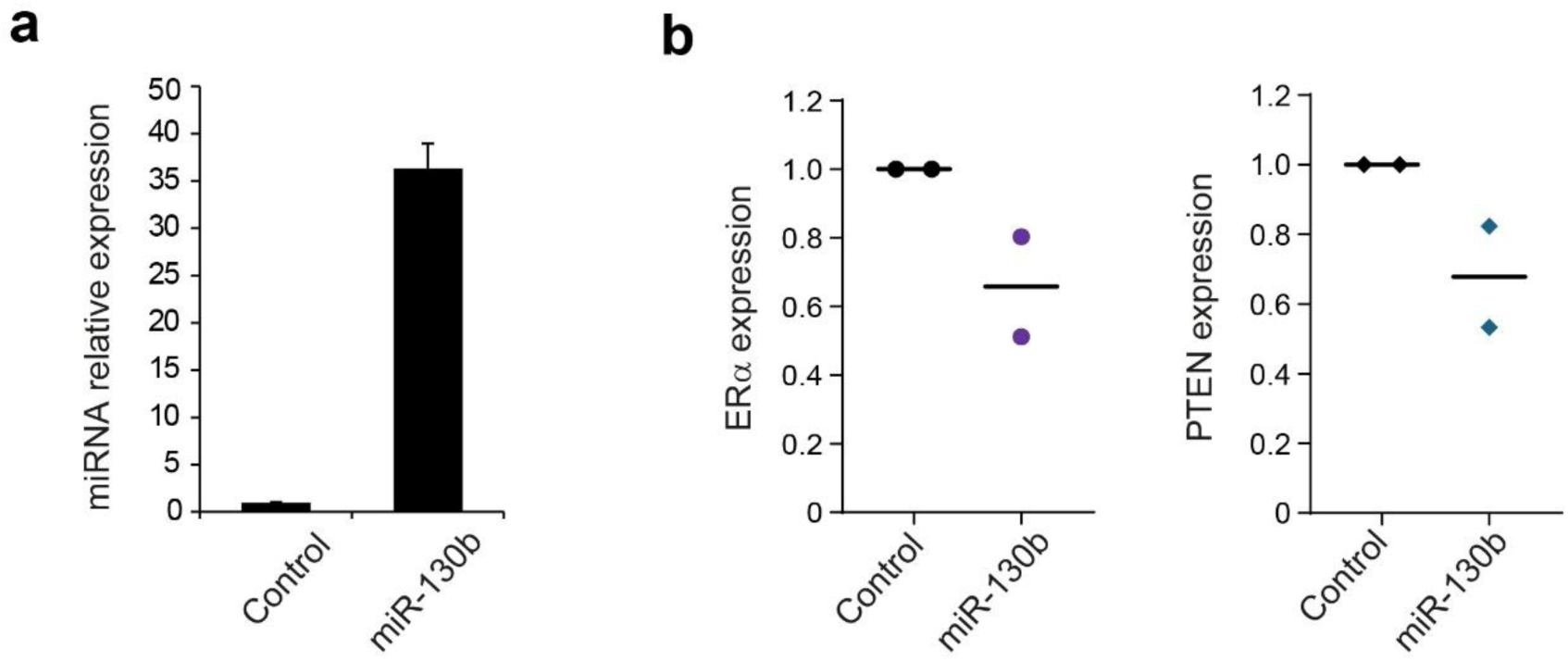
Regulation of ERα and PTEN by miR-130b. **a,** Bar graph showing relative expression of miR-130b in WEHI-control and WEHI-miR-130b cells determined by Taqman Assay. **b,** Graphs showing quantification of protein expression levels of ERα and PTEN in WEHI-control and WEHI-miR-130b cells determined by immunoblot in two independent experiments. The levels of these proteins were normalized to that of β-actin (loading control).

**Extended Data Fig. 3.**
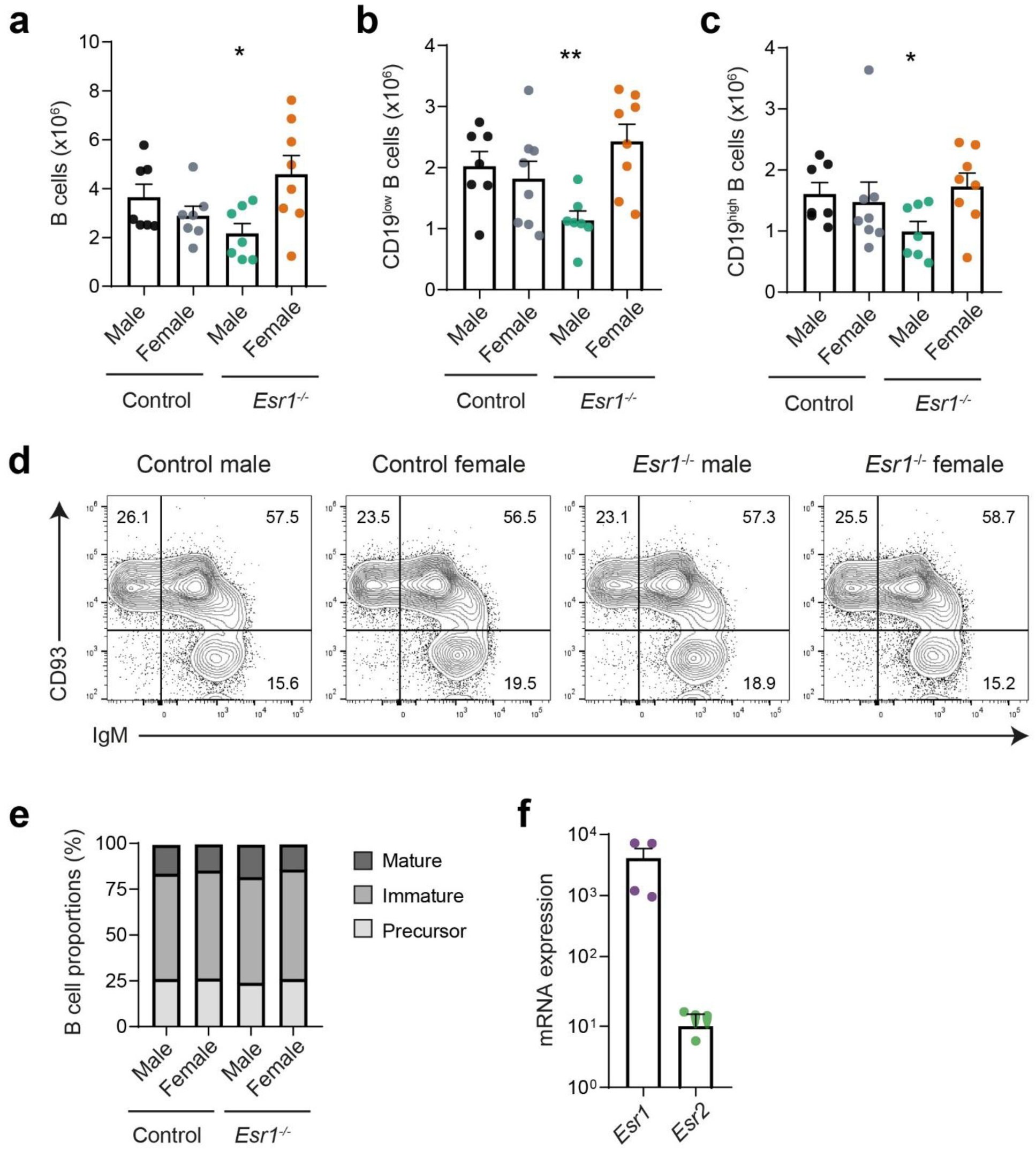
**a-c.** Bar graphs showing total numbers of CD19^+^-IgM^+^ (**a**), CD19^low^-IgM^+^ (**b**) and CD19^high^-IgM^+^ (**c**) cells in the bone marrow of control and *Esr*1^-/-^ male and female mice, as indicated. **d,** Representative contour plots showing percentages of immature (CD93^+^-IgM^+^), mature (CD93^-^-IgM^+^) and precursor (CD93^-^-IgM^-^) B cells within total bone marrow CD19^+^ B cells. **e,** Bar graph showing average percentages of immature, mature and precursor B cells within the total bone marrow B cell compartment. **f,** Relative mRNA levels of *Esr1* and *Esr2* in developing B cells in the bone marrow of female control mice. Graphs represent mean + s.e.m. Data are pooled from three independent experiments. n=7 mice (control and Esr1^-/-^ female groups), and n=8 mice (control and Esr1^-/-^ male groups). Statistical significance was determined with unpaired two-tailed Student’s t tests. * P ≤ 0.05 and ** P ≤ 0.01.

**Extended Data Fig. 4.**
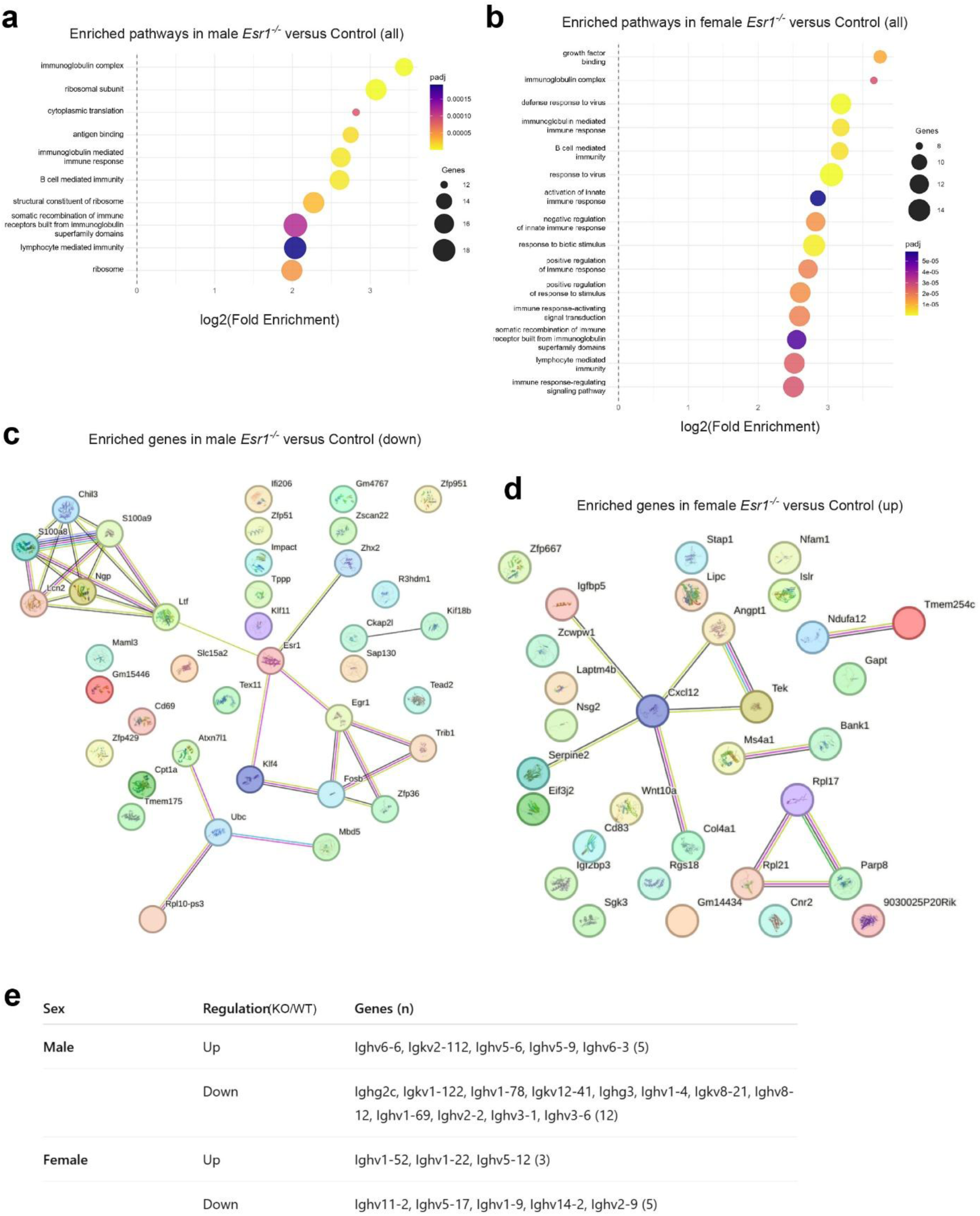
**a,b.** GO enrichment analysis showing pathways enriched in genes in developing B cells in the bone marrow of male *Esr1^-/-^* mice compared with their control counterparts (**a**), or of female *Esr1^-/-^* cells compared with the control group (**b**). **c,d,** STRING interactions analysis showing interconnections enriched in genes downregulated in developing B cells in the bone marrow of male *Esr1^-/-^* mice compared with their control counterparts (**c**), or upregulated in female *Esr1^-/-^* mice compared with the control group (**d**). **e,** Immunoglobulin gene segments upregulated or downregulated in male and female *Esr1*-deficient mice compared with their control counterparts, as indicated.

**Extended data Fig. 5.**
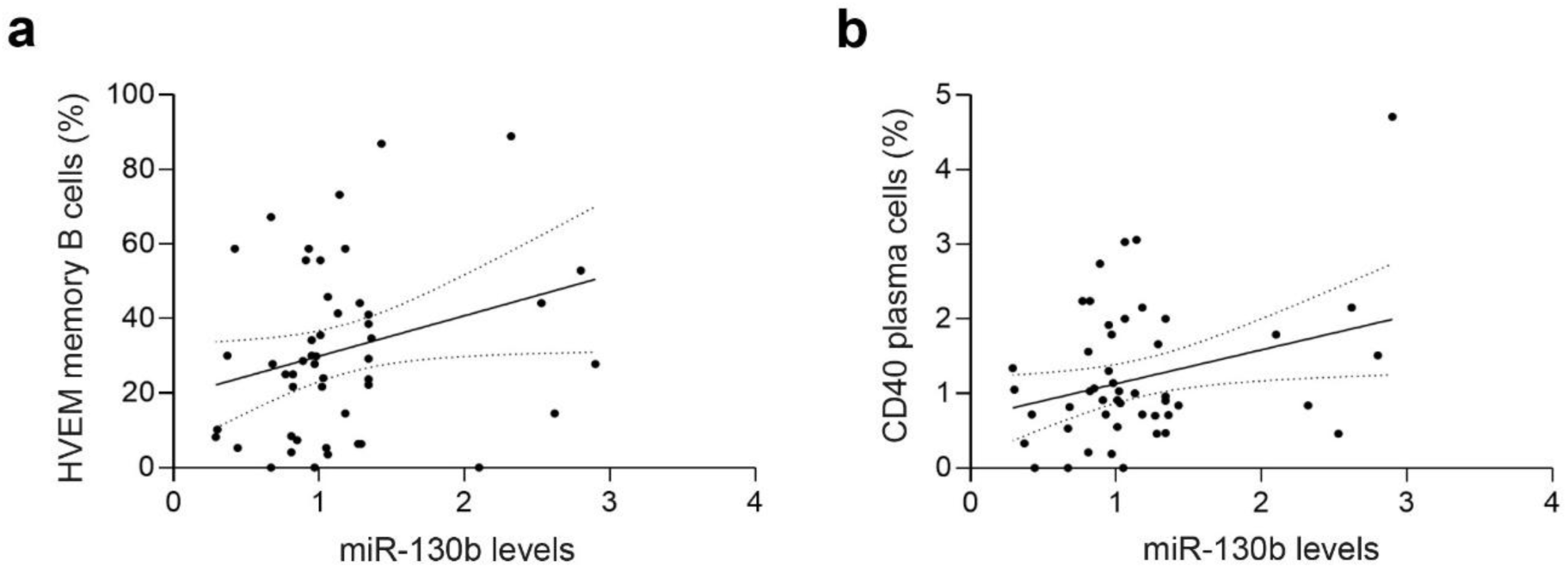
Elevated miR-130b in circulating extracellular vesicles correlates with increased circulating HVEM^+^ memory B cells as well as CD40 and PDL1^+^ plasma cells. **a-c,** Graphs showing correlation of miR-130b in circulating extracellular vesicles (EVs) of patients with multiple sclerosis with circulating HVEM^+^ memory B cells (**a**), and CD40^+^ plasma cells (**b**). n=50 patients. Statistical analysis was performed with a Spearman correlation test, P= 0.017 in **a**, P=0.01 in b, and P=0.031 in **c**.

**Extended Data Table 1.** List of lymphocyte-expressed miRNAs used for Multiple Em for Motif Elicitation (MEME) analysis.

| **Lymphocyte-expressed miRNA families** |  |  |
| --- | --- | --- |
| Let-7 | miR-194 | miR-342 |
| miR-1 | miR-1948 | miR-345 |
| miR-10 | miR-196 | miR-350 |
| miR-101 | miR-1964 | miR-351 |
| miR-103 | miR-1981 | miR-361 |
| miR-1198 | miR-199 | miR-362 |
| miR-122 | miR-203 | miR-374 |
| miR-127 | miR-204 | miR-375 |
| miR-128 | miR-21 | miR-378 |
| miR-130 | miR-210 | miR-423 |
| miR-132 | miR-219 | miR-425 |
| miR-138 | miR-22 | miR-450 |
| miR-139 | miR-221 | miR-451 |
| miR-140 | miR-223 | miR-455 |
| miR-142 | miR-23 | miR-504 |
| miR-143 | miR-24 | miR-505 |
| miR-146 | miR-26 | miR-511 |
| miR-147 | miR-27 | miR-542 |
| miR-148 | miR-28 | miR-574 |
| miR-15 | miR-29 | miR-598 |
| miR-150 | miR-296 | miR-652 |
| miR-155 | miR-298 | miR-671 |
| miR-17 | miR-30 | miR-672 |
| miR-181 | miR-31 | miR-674 |
| miR-184 | miR-32 | miR-676 |
| miR-185 | miR-324 | miR-7 |
| miR-186 | miR-326 | miR-700 |
| miR-188 | miR-328 | miR-744 |
| miR-19 | miR-33 | miR-8 |
| miR-190 | miR-330 | miR-872 |
| miR-191 | miR-331 | miR-92 |
| miR-192 | miR-339 | miR-95 |
| miR-193 | miR-34 | miR-96 |
| miR-1934 | miR-340 |  |

**Extended Data Table 2.**
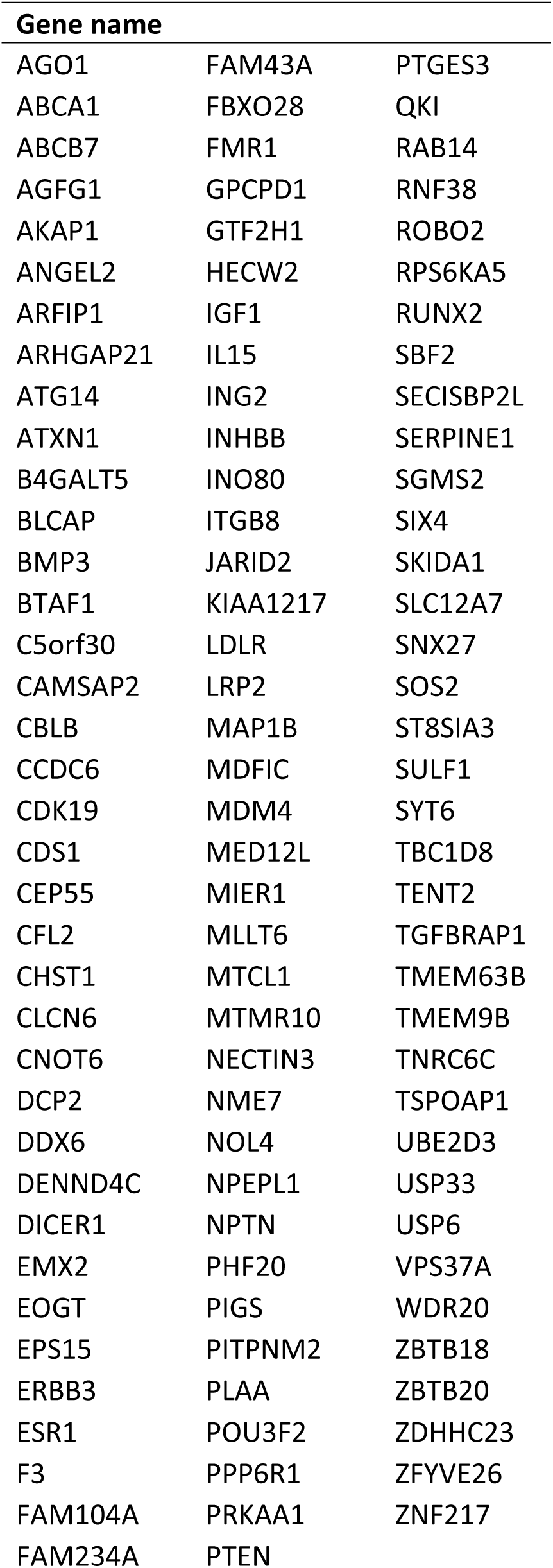
Predicted common target genes of miR-130b, miR-19b and miR-148a identified through RDB target mining.

**Extended data Table 3.**
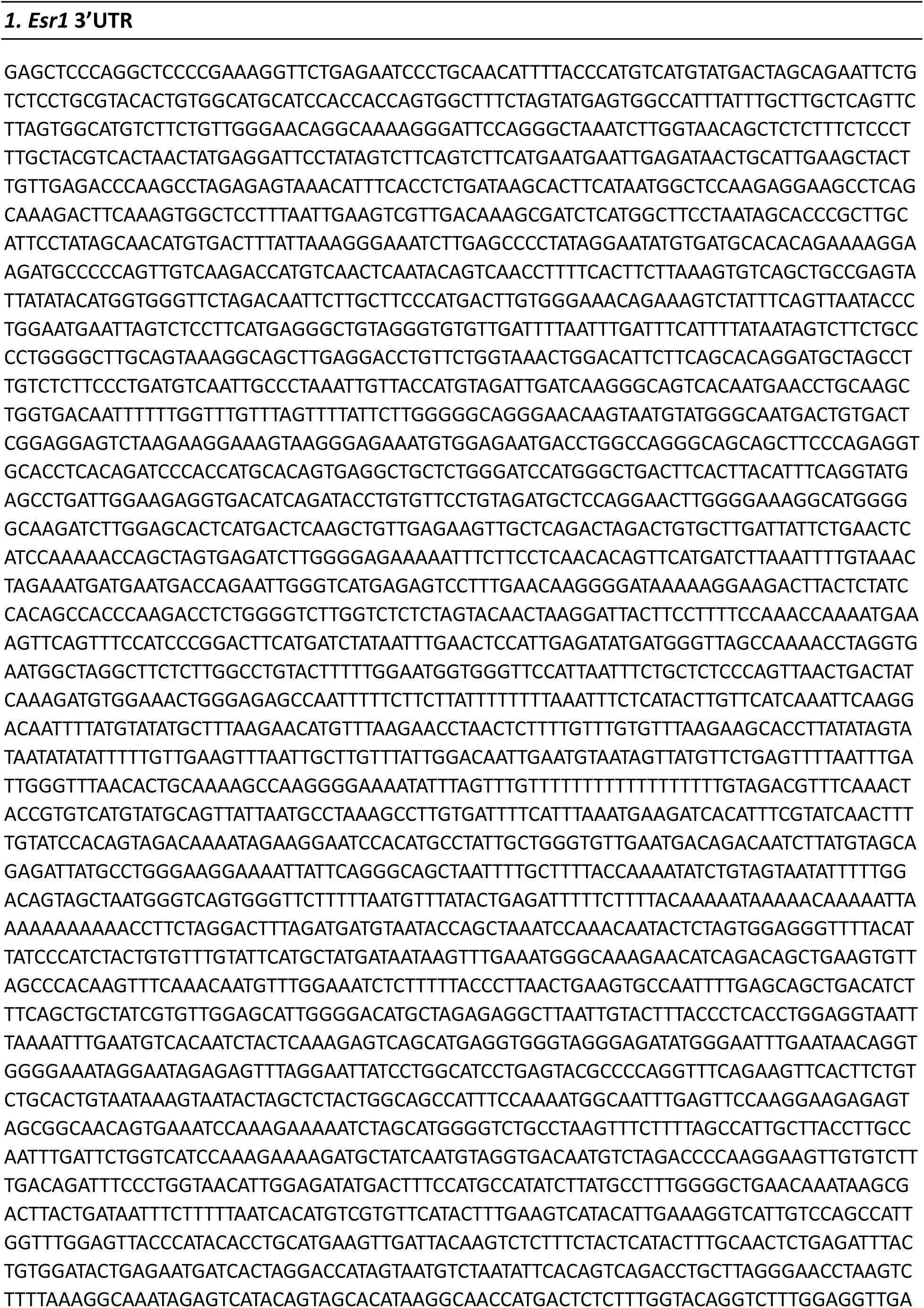
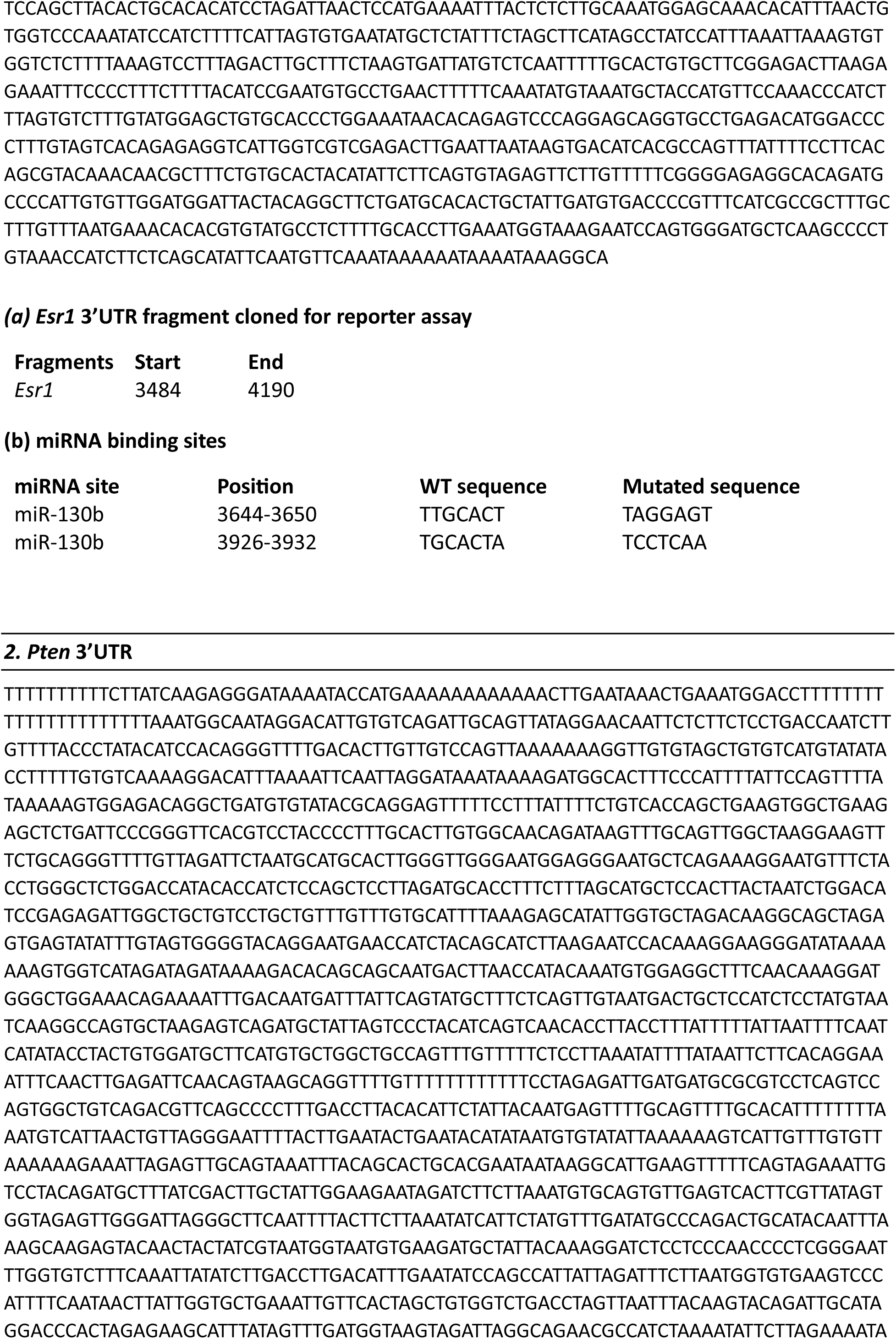
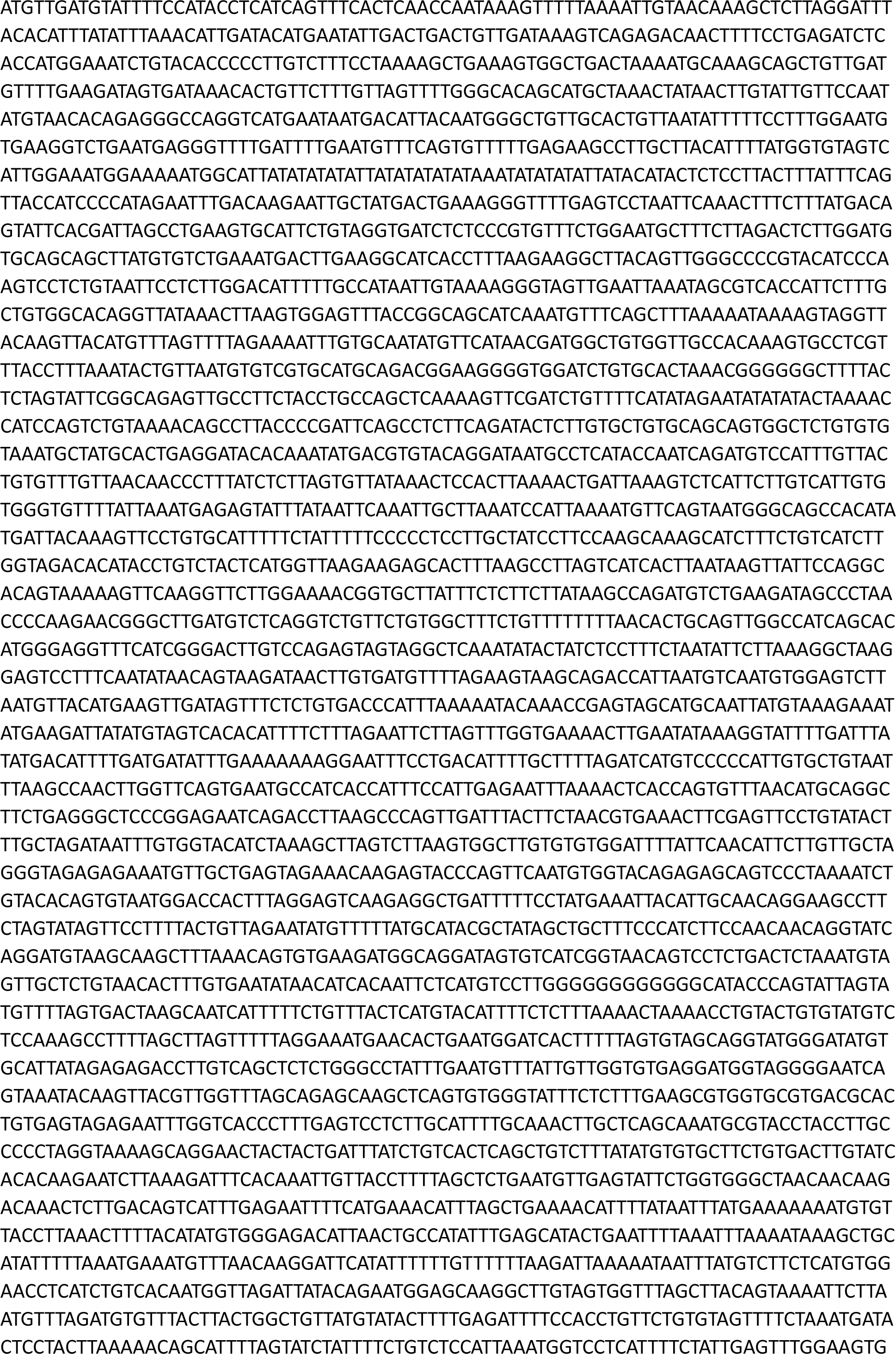
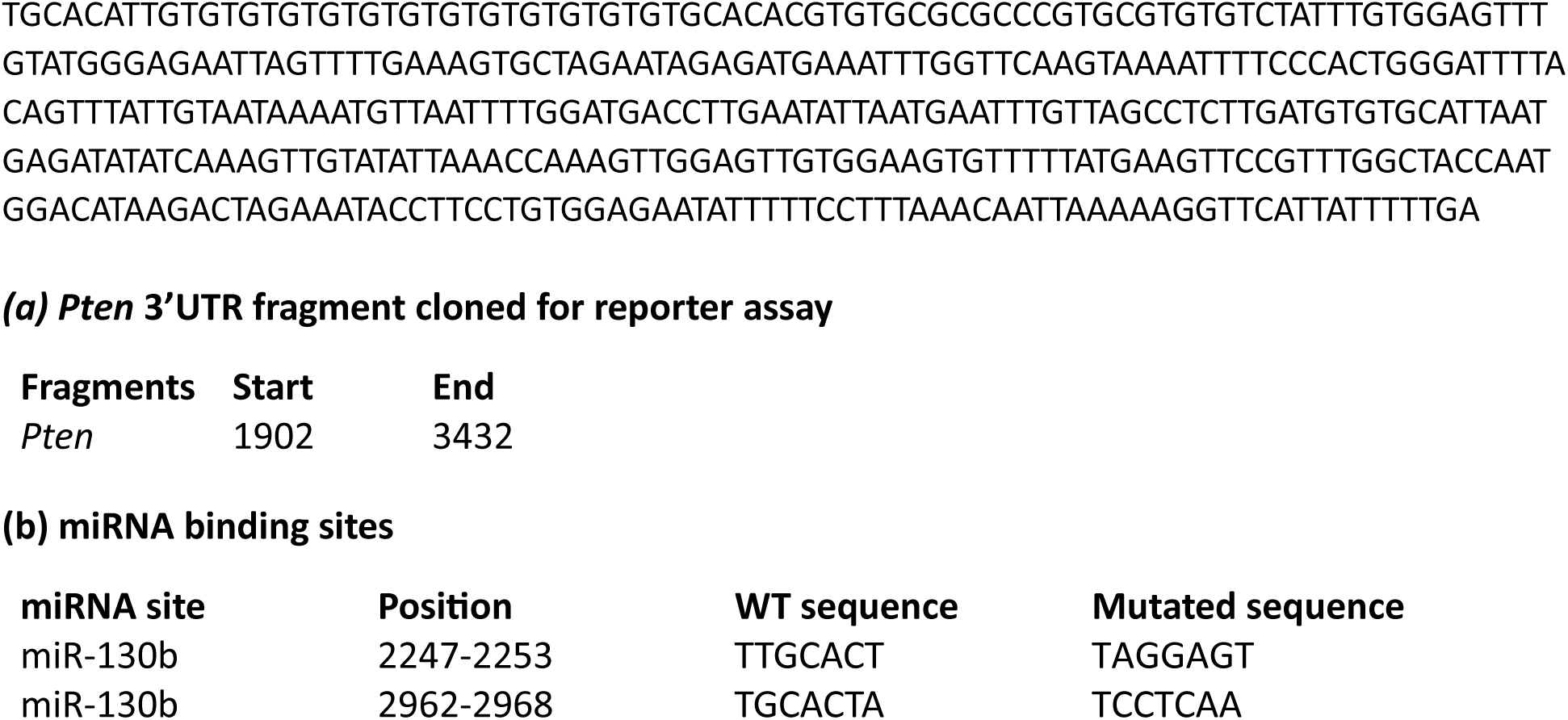
***Esr1* and *Pten* 3′ UTR cloned fragments for reporter assays, and mutated binding sites for miR-130b (*Esr1* and *Pten*).**

**Extended Data Table 4.** Demographic and clinical data of the patients with multiple sclerosis included in the study.

|  | **MS patients**<br>**(n=41)** |
| --- | --- |
| **Demographics** |  |
| Women. n (%) | 22 (53.7%) |
| Age (years) | 43.12 (9.43) |
| **Clinical data** |  |
| Time from diagnosis (months) | 96.03 (108.86) |
| Baseline SDMT | 47.40 (13.53) |
| Brain volume (cc) | 1119,85 (156,86) |
| Gray matter volume (cc) | 603,45 (72,07) |
| Cortical gray matter volume (cc) | 710,21 (1206,50) |
| Caudate nucleus volume (cc) | 6,68 (1,12) |
| Cerebellum volume (cc) | 7137,24 (15,97) |

\* All values represent mean (SD) unless otherwise indicated.
Abbreviations: CC, cubic centimeters; N, number; SD, standard deviation; SDMT, symbol digit modality test.

## Materials and Methods

### Mice

The generation of IgM^b^-macroself mice (MMRRC Strain 36510), *Pten*^fl/fl^ mice (Jax stock 006440), and *Esr1* knockout mice (Jax stock 026176) was previously described (*11, 34–36*). Mice were housed and bred in the animal facility of Institute for Biomedical Research, Autonoma University of Madrid under specific pathogen-free conditions. All animal experiments were approved by Animal Care and Bioethics Committee of Institute for Biomedical Research, Autonoma University of Madrid, and Community of Madrid, Spain.

### Retrovirus production

Retroviruses were generated by cotransfecting HEK293T cells with constructs encoding miR-130b, miR-33, miR-210 or an empty vector (control) from the pMIRWAY library, which includes green fluorescent protein (GFP) as a reporter, alongside GAG-POL and VSV-G plasmids. Confluent HEK293T cells were plated at a dilution of 1:5 the day before transfection. Cells were co-transfected with individual miRNAs or control plasmids, and packaging constructs. Cell culture medium was replaced after 24 hours. At 48 hours post-transfection, virus-containing supernatants were harvested, filtered, aliquoted, and stored at −80°C until use. Viral titers were determined by serial dilution (1:5) of the viral supernatants and analysis of GFP expression in infected target cells 48 hours after infection.

### Cell lines

WEHI-231 cells and their derivatives were grown in DMEM with 10% FBS, 2 mmol/l L-glutamine and Pen/Strep, and 50 μM 2-mercaptoethanol. The stable cell lines WEHI-control and WEHI–miR-130b were generated by retroviral transduction of WEHI-231 cells with particles encoding miR-130b or an empty construct (control), and enrichment with FACS. Specifically, WEHI-231 cells at 0.5×10^6^ cells/ml were transduced with MSCV-control-GFP or MSCV-miR-130b-GFP retroviral particles in the presence of 8 μg/ml polybrene. Cells were expanded for 48 hours, sorted by FACS to enrich the GFP^+^ infected population, grown for an additional 48 hours, aliquoted and stored in liquid nitrogen until use.

### Bone marrow transduction and reconstitution experiments

Donor 6-8 weeks-old C57BL/6J mice were injected with 5-fluorouracil (Sigma) at a concentration of 150 mg/kg. After 4 days, bone marrow cells were isolated and HSPCs were enriched by Ficoll (GE Healthcare) density-gradient centrifugation. Cells were resuspended in X-VIVO 15 medium (Lonza) supplemented with 10% BSA, 2 mmol/l L-glutamine and Pen/Strep, 100 μM 2-mercaptoethanol, 100 ng/ml mSCF, 50 ng/ml mTPO, 50 ng/ml mFlt3L and 20 ng/ml mIL-3, and transduced overnight with retroviral particles on retronectin-coated plates (Takara) in the presence of 2 μg/ml polybrene (Sigma Merk). Following infection, cells were retrieved, washed with excess PBS, and injected into the tail vein of recipient 8-12 weeks-old IgM^b^-macroself mice that had been previously irradiated (on the same day) with two doses of 5.0 Gy separated by 3 hours. Both female and male recipient mice were used. Each recipient mouse received 2×10^6^ to 5×10^6^ infected cells mixed with 0.5×10^6^ helper bone marrow cells from CD45.1^+^ C57BL/6J mice. 8 weeks after reconstitution, recipient mice were euthanized and the presence of B cells in the spleen was determined by cell surface staining with anti-CD19-PE (6D5; BioLegend) and anti-IgM-Cy5 (115-175-075; Jackson ImmunoResearch), and flow cytometry analysis in a FACS Canto II (BD Biosciences). Data were analyzed using the FlowJo v10 software.

### Purification of murine primary immature B cells

Femur and tibia bones were isolated from C57BL/6J mice, and their bone marrow cells retrieved by flushing in PBS 2% FBS. Cells were washed twice with PBS 2% FBS and enriched for B cells using magnetic beads (Miltenyi Biotec). Cells were then stained with anti-IgM-Cy5 (115-175-075; Jackson ImmunoResearch), CD19-BV421 (1D3; BD Biosciences) and CD93-PeCy7 (AA4.1; eBioscience). Immature B cells, defined as CD19^+^-IgM^+^-CD93^+^ cells, were sorted in a Moflow ASTRIOS Sorter (Beckman Coulter). Purified immature B cells were washed once with PBS and stored at −80 °C until use.

### Analysis of B cell development

Total cells were obtained from femur and tibia of male and female control and Esr1^-/-^ mice, erythrocytes were lysed with ACK lysis buffer and counted. Cells were stained with anti-IgM (115-175-075; Jackson ImmunoResearch), CD19 (1D3; BD Biosciences) and CD93 (AA4.1; eBioscience) coupled to fluorophores and analyzed by flow cytometry. Developing B cells were defined as CD19^+^-IgM^+^-CD93^+^ (immature) and CD19^+^-IgM^-^-CD93^+^ (precursor) cells. Mature cells were defined as CD19^+^-IgM^+^-CD93^-^ cells. Data were acquired in a FACS Canto II (Beckman Dickinson) and a Cytoflex S (Beckman Coulter) and analyzed with the FlowJo v10 software.

### Quantitative RT-PCR

WEHI-control and WEHI–miR-130b cells were stimulated with 2 μg/ml anti-IgM (115-175-075; Jackson ImmunoResearch) for 14 hours, and total RNA was extracted from the cells using TRI Reagent (Invitrogen), following the manufacturer’s instructions. Total RNA from purified mouse immature B cells was prepared using the RNeasy Micro Kit (Qiagen). cDNA was synthesized by reverse transcription of total RNA (1 μg) with the High-capacity cDNA reverse transcription kit (Applied Biosystems). Quantitative RT-PCR was performed using the Power SYBR Green PCR Master Mix System (Applied Biosystems) with specific primers for mouse *Cfl2, Esr1, Prkaa1, Pten, Runx2* and *Sos2*, using *Actb* for normalization. The following primers were used for quantitative RT-PCR analysis: *Cfl2* 5′-GCATCTGGAGTTACAGTGAATGA-3′ and 5′-CACCAATGTCACCCACCAAGA-3′; *Esr1* 5′-CCCGCCTTCTACAGGTCTAAT-3′ and 5′-CTTTCTCGTTACTGCTGGACAG-3′; *Igf1* 5′-CTGGACCAGAGACCCTTTGC-3′ and 5′-GGACGGGGACTTCTGAGTCTT-3′; *Prkaa1* 5′-GTCAAAGCCGACCCAATGATA-3′ and 5′-CGTACACGCAAATAATAGGG GTT-3′; *Pten* 5′-TGGGGAAGTAAGGACCAGAG-3′ and 5′-GGCAGACCACAAACTGA GGA-3′; *Runx2* 5′-AAAGCCAGAGTGGACCCTTCCA-3′ and 5′-ATAGCGTGCTGCCATT CGAGGT-3′; *Sos2* 5′-CAAGATGTTGAGGAACGAGTTCA-3′ and 5′-TGTCCACAGGTA GTAAGAGAGGA-3′; and *Actb* 5′-CTAAGGCCAACCGTGAAAG-3′ and 5′-ACCAG AGGCATACAGGGACA-3′.

### Transcriptome analysis

Bone marrow developing B cells were purified from femur and tibia of male and female control and Esr1^-/-^ mice using magnetic negative selection with the mouse B cell isolation kit (Miltenyi) followed by positive selection with CD93-microbeads (Miltenyi). Total RNA from these samples were obtained with the RNeasy Plus Mini Kit (Quiagen) following the manufactureŕs instructions. Messenger RNA was then purified from total RNA using poly-T oligo-attached magnetic beads. After fragmentation, the first strand cDNA was synthesized using random hexamer primers followed by the second strand cDNA synthesis. The library was ready after end repair, A-tailing, adapter ligation, size selection, amplification, and purification. The kit for library preparation was the Novogene NGS RNA Library Prep Set (PT042).

The library was checked with Qubit 2.0, real-time PCR for quantification, and a bioanalyzer 2100 for size distribution detection. Quantified libraries were pooled and sequenced on the Illumina platform NovaseqX plus, according to effective library concentration and data amount (9 Gb). The sequencing strategy was Pair-end 150 bp (PE150). The index-coded samples were clustered according to the manufacturer’s instructions. After cluster generation, the library preparations were sequenced on the Illumina platform Novaseq X plus, and paired-end reads were generated.

### Western blot

Dry cell pellets were resuspended in RIPA lysis buffer (150 mM NaCl, 1% NP-40, 0.5% Sodium deoxycholate, 0.1% SDS, 25 mM Tris pH 7.4, and ddH_2_O) supplemented with protease inhibitors (Roche) and phosphatase inhibitors (Sigma-Aldrich). Protein quantification was performed using the BCA assay (ThermoScientific). Protein extracts (10 μg per sample) were resolved on 10% SDS-PAGE gels and transferred onto nitrocellulose membranes (GE Healthcare). Membranes were blocked in TBS-0.1% Tween-20 (TBS-T) containing 5% BSA for one hour at room temperature, followed by washing with TBS-T. The membranes were then incubated overnight at 4°C with primary antibodies specific for ERα (Santa Cruz Biotechnology, sc-542), PTEN (Cell Signaling, 138G6), and β-actin (Santa Cruz Biotechnology, sc-47778). After extensive washing with TBS-T, membranes were incubated for one hour at room temperature with goat anti-rabbit or goat anti-mouse horseradish peroxidase-conjugated secondary antibodies (Santa Cruz Biotechnology). Bound antibodies were detected using the Clarity Western ECL Substrate (Bio-Rad) and visualized using a Fusion Solo S (Vilber) chemiluminescence imaging system.

### Dual luciferase reporter assays

The sequence encoding miR-130b was cloned into the expression vector pCXN2, and 3′ UTR fragments of *Esr1* and *Pten* were cloned into the psiCHECK2 vector (Promega). The miR-130b binding sites in the 3′ UTR of each target gene were mutated with the QuikChange Multi Site-Directed Mutagenesis Kit (Agilent Technologies). The resulting pCXN2 and psiCHECK2 plasmids were transfected into HEK293 cells using Fugene HD (Promega) according to the manufacturer’s instructions. After 24 hours, cells were lysed, and luciferase expression was measured using the Dual-luciferase assay system (Promega) following the manufacturer’s protocol. The renilla luciferase activity was normalized to that of firefly luciferase, and the ratio of renilla luciferase activity to firefly luciferase activity was arbitrarily set as 1 for the control pCXN2 vector for each psiCHECK2 reporter.

### Bone marrow transplantation

Donor bone marrow was harvested from 6-8 weeks-old mice of indicated genotypes and used to reconstitute lethally irradiated 8-12 weeks-old IgM^b^-macroself mice. For each recipient mouse, 2×10^6^ to 5×10^6^ donor bone marrow cells were mixed with 0.25×10^6^ helper bone marrow cells from CD45.1^+^ C57BL/6J mice and were inoculated intravenously into IgM^b^-macroself mice that had been previously irradiated (on the same day) with two doses of 5 Gy separated by 3 hours. Both female and male recipient mice were used unless otherwise stated. Splenocytes were analyzed 8 weeks after reconstitution to determine the presence of B cells by staining with anti-CD19-PE (6D5; BioLegend) and anti-IgM-Cy5 (115-175-075; Jackson ImmunoResearch) and flow cytometry analysis as above.

### Determination of miR-130b levels in EVs of patients with multiple sclerosis

An observational, longitudinal, and prospective study was conducted, including patients with a confirmed diagnosis of relapsing-remitting multiple sclerosis (RRMS), based on the McDonald criteria, from May 2021 to June 2023. Only patients not receiving immunomodulatory treatment were included. The study was approved by the Research Ethics Committee of La Paz University Hospital (PI-4675), and informed consent was obtained from each participant. Demographic and clinical data collected included sex, age, disease duration, Symbol Digit Modalities Test (SDMT) scores, new white matter lesions on T2-weighted MRI sequences, total brain volume, and the volumes of specific structures, including white matter, gray matter, cerebellum, hypothalamus, and basal ganglia, assessed using 3D T1 and FLAIR sequences and analyzed with aCloud and FreeSurfer v5.3.0.

Blood samples were collected in 9 mL EDTA tubes and centrifuged at 1800 rpm for 15 minutes to isolate cell-free plasma. For extracellular vesicle (EV) isolation, 1 mL of plasma per patient was thawed rapidly in a 37°C water bath, transferred to a 1.5 mL Eppendorf tube and centrifuged at room temperature for 20 minutes at 2000g. EVs were isolated using the ExoQuick EV isolation kit (System Biosciences, USA). miR-130b content in EVs was analyzed using miRNA sequencing. Total RNA was extracted from EVs with the miRNeasy Serum/Plasma Advanced Kit (Qiagen, USA). Library preparation was performed from 10 ng of total RNA per sample with the SMARTer smRNA-Seq Kit (Illumina, USA) and validated for size, purity, and concentration using the Agilent Bioanalyzer (Agilent Technologies, USA). Samples were pooled in equimolar amounts and sequenced on the Illumina HiSeq 2500 platform to generate 101 base pair reads. Read quality was assessed with FastQC (Babraham Bioinformatics, UK), and sequences were trimmed and quality-filtered with Cutadapt v4.7. High-quality reads were aligned to miRNA sequences from miRBase v22 using the mirDeep2 tool. On the other hand, peripheral blood samples were collected from 54 healthy controls and 64 patients with multiple sclerosis (MS). Circulating extracellular vesicles were isolated, and total RNA was extracted using the exoRNeasy Mini Kit (Qiagen) according to the manufacturer’s instructions. Quantification of miR-130b was performed using the TaqMan™ Advanced miRNA cDNA Synthesis Kit followed by quantitative PCR using the Applied Biosystems 7300 Real-Time PCR System. miRNA-16 was used as the endogenous control for normalization of relative expression levels. All reactions were performed in duplicate.

### Bioinformatic analyses

For Multiple Em for Motif Elicitation analysis, a list of 101 lymphocyte-expressed miRNA families was generated from available expression profiling data (*33*) and the nucleotides in positions 1-8 filtered as a FASTA file. This was submitted as input for analysis in Multiple Em for Motif Elicitation (MEME) suite (https://meme-suite.org/meme/tools/meme) as previously described (*37*). The output was ranked by MEME based on the E-value obtained for each motif.

To identify protein-coding genes with potential functions in B cell tolerance, the predicted target genes of miR-130b, miR-19b and miR-148a were determined individually through RDB miRNA mining (*38*). The common targets were then selected, and the resulting 110 genes subjected to Ingenuity Pathway Analysis (IPA, Qiagen) for identification of potential functionally relevant signaling pathways regulated by these miRNAs.

### Statistical analysis

Data were analyzed using an unpaired two-tailed Student’s t test. Statistical significance was defined as * P ≤ 0.05, ** P ≤ 0.01, *** P ≤ 0.001 and **** P ≤ 0.0001. Bar graphs show mean + s.e.m or mean + s.d., as indicated for each Figure.

## Data availability

All data included in this manuscript are available from the corresponding author, A.G-M., upon reasonable request.

## Acknowledgments

We thank Luis del Peso for his help with the MEME analysis. We also thank the personnel from the animal facilities at IIBM and UAM, and from UAM Flow Cytometry Core for technical assistance. This work was supported by the grants RTI2018-100008-A-I00 MCIN/AEI/10.13039/501100011033 and FEDER, PID2021-128244OB-I00/AEI/FEDER 10.13039/501100011033 and CNS2022-136069/MCIN/AEI/10.13039/501100011033 and European Union “NextGenerationEU”/PRTR from Spanish Ministry of Science and Innovation, SI1-PJI-2019-00241 from Community of Madrid, LABAE20001GONZ from Spanish Association Against Cancer (AECC), XXIII Beca FERO from FERO Foundation, and XXI National Research Grants in Life Sciences from Ramon Areces Foundation to A.G-M. A.G-M was supported by merit award RyC-21155 MCIN/AEI/10.13039/ 501100011033 and FSE and L.G-R by the PRE2019-087940 FPI fellowship from Spanish Ministry of Science and Innovation. MdP.G-M. was supported by an FPI-UAM 2020 fellowship from Autonoma University of Madrid and L.M. by the PEJ-2020-AI/BMD-17617 contract from the Garantia Juvenil Program of Community of Madrid.

## Author contributions

A.G-M. conceptualized and designed the project and wrote the manuscript with contributions from all authors. JL.D-V., B.H-F., MdP.G-M., AM.P-M., J.S-G., M.M-H., T.G-S., A.M., and A.G-M. performed experiments. L.G-R. performed bioinformatic analyses. MP.L-M and L.O. performed all experiments with samples of patients with autoimmunity and provided materials and reagents. L.M. managed and genotyped the mouse colonies, and provided technical support. D.N. and C.X. supported experiments and provided materials and reagents. A.G-M. obtained funding and supervised the project.

## References

1. E. Zerhouni. (Progress in autoimmune diseases research. N.I.o.H. U.S. Department of Health and Human Services, ed. , 2005).

2. N. Conrad et al., Incidence, prevalence, and co-occurrence of autoimmune disorders over time and by age, sex, and socioeconomic status: a population-based cohort study of 22 million individuals in the UK. Lancet 401, 1878–1890 (2023).

3. F. W. Miller, The increasing prevalence of autoimmunity and autoimmune diseases: an urgent call to action for improved understanding, diagnosis, treatment, and prevention. Curr Opin Immunol 80, 102266 (2023).

4. K. S. Forsyth, N. Jiwrajka, C. D. Lovell, N. E. Toothacre, M. C. Anguera, The conneXion between sex and immune responses. Nat Rev Immunol, (2024).

5. M. J. Shlomchik, Sites and stages of autoreactive B cell activation and regulation. Immunity 28, 18–28 (2008).

6. S. Yurasov et al., Defective B cell tolerance checkpoints in systemic lupus erythematosus. J Exp Med 201, 703–711 (2005).

7. L. Menard, J. Samuels, Y. S. Ng, E. Meffre, Inflammation-independent defective early B cell tolerance checkpoints in rheumatoid arthritis. Arthritis Rheum 63, 1237–1245 (2011).

8. T. Kinnunen et al., Specific peripheral B cell tolerance defects in patients with multiple sclerosis. J Clin Invest 123, 2737–2741 (2013).

9. D. Nemazee, Mechanisms of central tolerance for B cells. Nat Rev Immunol 17, 281–294 (2017).

10. H. Wardemann, M. C. Nussenzweig, B-cell self-tolerance in humans. Adv Immunol 95, 83–110 (2007).

11. B. H. Duong et al., Negative selection by IgM superantigen defines a B cell central tolerance compartment and reveals mutations allowing escape. J Immunol 187, 5596–5605 (2011).

12. A. Gonzalez-Martin et al., The microRNA miR-148a functions as a critical regulator of B cell tolerance and autoimmunity. Nat Immunol 17, 433–440 (2016).

13. M. Lai et al., Regulation of B-cell development and tolerance by different members of the miR-17∼92 family microRNAs. Nat Commun 7, 12207 (2016).

14. D. P. Bartel, MicroRNAs: target recognition and regulatory functions. Cell 136, 215–233 (2009).

15. C. Xiao, K. Rajewsky, MicroRNA control in the immune system: basic principles. Cell 136, 26–36 (2009).

16. K. M. Pauley, S. Cha, E. K. Chan, MicroRNA in autoimmunity and autoimmune diseases. J Autoimmun 32, 189–194 (2009).

17. W. Wang et al., Up-regulation of Serum MiR-130b-3p Level is Associated with Renal Damage in Early Lupus Nephritis. Sci Rep 5, 12644 (2015).

18. C. D. Browne, C. J. Del Nagro, M. H. Cato, H. S. Dengler, R. C. Rickert, Suppression of phosphatidylinositol 3,4,5-trisphosphate production is a key determinant of B cell anergy. Immunity 31, 749–760 (2009).

19. S. Sato, N. Ono, D. A. Steeber, D. S. Pisetsky, T. F. Tedder, CD19 regulates B lymphocyte signaling thresholds critical for the development of B-1 lineage cells and autoimmunity. J Immunol 157, 4371–4378 (1996).

20. H. D. Phan et al., CD24 and IgM Stimulation of B Cells Triggers Transfer of Functional B Cell Receptor to B Cell Recipients Via Extracellular Vesicles. J Immunol 207, 3004–3015 (2021).

21. D. C. Ayre et al., Dynamic regulation of CD24 expression and release of CD24-containing microvesicles in immature B cells in response to CD24 engagement. Immunology 146, 217–233 (2015).

22. S. Ghosh, R. S. Klein, Sex Drives Dimorphic Immune Responses to Viral Infections. J Immunol 198, 1782–1790 (2017).

23. S. L. Klein, K. L. Flanagan, Sex differences in immune responses. Nat Rev Immunol 16, 626–638 (2016).

24. M. K. Desai, R. D. Brinton, Autoimmune Disease in Women: Endocrine Transition and Risk Across the Lifespan. Front Endocrinol (Lausanne) 10, 265 (2019).

25. T. S. Thurmond et al., Role of estrogen receptor alpha in hematopoietic stem cell development and B lymphocyte maturation in the male mouse. Endocrinology 141, 2309–2318 (2000).

26. G. J. Shim, L. L. Kis, M. Warner, J. A. Gustafsson, Autoimmune glomerulonephritis with spontaneous formation of splenic germinal centers in mice lacking the estrogen receptor alpha gene. Proc Natl Acad Sci U S A 101, 1720–1724 (2004).

27. D. H. Kim et al., Estrogen receptor α in T cells suppresses follicular helper T cell responses and prevents autoimmunity. Exp Mol Med 51, 1–9 (2019).

28. V. Rider et al., Gender Bias in Human Systemic Lupus Erythematosus: A Problem of Steroid Receptor Action? Front Immunol 9, 611 (2018).

29. A. Di Cristofano et al., Impaired Fas response and autoimmunity in Pten+/- mice. Science 285, 2122–2125 (1999).

30. G. L. Mutter, M. C. Lin, J. T. Fitzgerald, J. B. Kum, C. Eng, Changes in endometrial PTEN expression throughout the human menstrual cycle. J Clin Endocrinol Metab 85, 2334–2338 (2000).

31. M. M. Scully, L. K. Palacios-Helgeson, L. S. Wah, T. A. Jackson, Rapid estrogen signaling negatively regulates PTEN activity through phosphorylation in endometrial cancer cells. Horm Cancer 5, 218–231 (2014).

32. K. Xu et al., Extracellular vesicles as potential biomarkers and therapeutic approaches in autoimmune diseases. J Transl Med 18, 432 (2020).

33. S. Kuchen et al., Regulation of microRNA expression and abundance during lymphopoiesis. Immunity 32, 828–839 (2010).

34. R. Lesche et al., Cre/loxP-mediated inactivation of the murine Pten tumor suppressor gene. Genesis 32, 148–149 (2002).

35. R. C. Rickert, J. Roes, K. Rajewsky, B lymphocyte-specific, Cre-mediated mutagenesis in mice. Nucleic Acids Res 25, 1317–1318 (1997).

36. S. C. Hewitt et al., Biological and biochemical consequences of global deletion of exon 3 from the ER alpha gene. FASEB J 24, 4660–4667 (2010).

37. T. L. Bailey, C. Elkan, Fitting a mixture model by expectation maximization to discover motifs in biopolymers. Proc Int Conf Intell Syst Mol Biol 2, 28–36 (1994).

38. Y. Chen, X. Wang, miRDB: an online database for prediction of functional microRNA targets. Nucleic Acids Res 48, D127–D131 (2020).

